# Clade V MLO proteins are bona fide host susceptibility factors required for powdery mildew pathogenesis in Arabidopsis

**DOI:** 10.1101/2025.06.18.660284

**Authors:** David Bloodgood, Qiong Zhang, Pai Li, Ying Wu, Michael Pan, Christina Zhou, Apsen Hsu, Jun Zhang, Ralph Panstruga, Sharon Kessler, Ping He, Libo Shan, Chang-I Wei, Shunyuan Xiao

## Abstract

Obligate biotrophic powdery mildew (PM) fungi strictly require living host to survive. To search for host factors or processes essential for PM pathogenesis, a tailored genetic screen was conducted with the immuno-compromised *eds1-2/pad4-1/sid2-2* (*eps*) triple Arabidopsis mutant. This led to the identification of five allelic disruptive mutations in *Mildew Locus O 2* (*MLO2*) to be responsible for the compromised-immunity-yet-poor infection (cipi) mutant phenotype upon challenge from an adapted PM isolate. Moreover, the *eds1/pad4/sid2/mlo2/mlo6/mlo12* (*eps3m*) sextuple mutant display near complete immunity to the adapted PM fungus without sign of defense activation, demonstrating that these three clade V MLOs in Arabidopsis are *bona fide* host susceptibility factors of PM fungi. Confocal imaging revealed focal accumulation of MLO2-GFP in the peri-penetration peg membranous space, implicating MLO2 in repairing and stabilizing the damaged host plasma membrane, which may be co-opted by PM fungi for haustorium differentiation. Results from domain-swapping analysis between MLO1 and MLO2 suggest a bipartite functional configuration for MLO2: its C-terminus determines where and when MLO2 functions, while its N-terminal seven transmembrane domain region executes the cellular function that is critical for PM pathogenesis. Genetic studies further demonstrate that, unlike MLO7 in synergids, focal accumulation of MLO2 does not depend on FERONIA (FER) and its five other family members, nor does it require phosphatidylinositol 4,5-bisphosphate produced from phosphatidylinositol 4-phosphate 5-kinase 1 (PIP5K1) and PIP5K2. Together, these findings define clade V MLOs as host factors co-opted by obligate biotrophic PM fungi for successful host colonization.

## Introduction

Powdery mildew (PM) fungi are obligate biotrophic pathogens that strictly require living host cells to survive and thrive (Schulze-Lefert and Panstruga, 2003). Upon landing on a host leaf surface, a PM spore germinates within six hours, forming an appressorium to penetrate the host cell wall within 7–10 hours. The sporeling then differentiates the haustorium physically inside the host cell from the tip of the penetration peg in 12-14 hours (Koh et al., 2005). Concomitant with the development of the haustorium, a host-derived extra-haustorial membrane (EHM) is formed to encase the haustorium (Wang et al., 2009; Berkey et al., 2017). The haustorium is believed to mature in 20-24 hours and then it is capable of extracting water and nutrient from the host cell to support hyphal growth on the leaf surface. Within four to five days, the fungus develops an extensive mycelial network, capable of forming conidiophores that produce new conidia, thereby completing its asexual life cycle. Like other obligate biotrophic pathogens, PM fungi cannot be cultured and are genetically intractable. Despite extensive studies into plant-PM fungal interactions, a critical question regarding PM biotrophy remains unresolved: aside from host-derived nutrients, are there any specific host factors or processes that are truly indispensable for PM pathogenesis? Conventional genetic screens have been conducted to identify mutants that show enhanced resistance to PM fungi. Many such mutants show lesion mimic or autoimmunity, with disruptive mutations in likely negative regulators of immunity (Frye and Innes, 1998; Tang et al., 2005, 2006; Wang et al., 2008; Zhang et al., 2008). To identify possible host factors or processes that are critical for PM pathogenesis without the complication of autoimmunity, we constructed a triple mutant in which genes encoding two essential immune signaling components, EDS1 (Falk et al., 1999) and PAD4 (Jirage et al., 1999), and a key enzyme, SID2, required for salicylic acid (SA) biosynthesis (Wildermuth et al., 2001) were mutated. This *eds1-2/pad4-1/sid2-2* (*eps*) mutant is super-susceptible to the adapted PM isolate *Golovinomyces cichoracearum* (*Gc*) UCSC1 (Zhang et al., 2018). We then performed a large-scale genetic screen using EMS-mutagenized seeds of *eps* to identify mutants that show **<u>c</u>**ompromised **i**mmunity yet **<u>p</u>**oor **i**nfection (*cipi*), with the goal of identifying potential host susceptibility factors required for PM pathogenesis. Among the 18 *cipi* mutants isolated, five mutants with the poorest infection each contains a causal mutation in *MLO2* (At1g11310) (see the Results section), suggesting that *mlo2*-conditioned poor infection can be uncoupled from defense activation.

The MILDEW LOCUS O (MLO) was originally discovered to confer broad-spectrum resistance to PM fungal isolates in barley (*Hordeum vulgare; Hv*) and recessive mutations in the *HvMLO* gene encoding a seven transmembrane-domain (7TM) protein are responsible for the resistance (Buschges et al., 1997). HvMLO belongs to a plant lineage-specific 7TM protein family (Kusch et al., 2016). Impairment of *Arabidopsis thaliana* (*At*) *MLO2*, *MLO6* and *MLO12* also results in near complete resistance to an adapted PM isolate (Consonni et al., 2006). These earlier findings stimulated targeted mutagenesis of *MLO* genes as a promising new avenue for creating PM-resistant crops in recent years (Feechan et al., 2008; Pavan et al., 2011; Zheng et al., 2013; Wang et al., 2014; Kusch and Panstruga, 2017; Nekrasov et al., 2017; Wan et al., 2020). Unfortunately, *mlo* mutants with strong or near-complete resistance to PM fungi often display reduced plant stature, elevated levels of SA, increased callose deposition, and early leaf senescence, resembling characteristics of autoimmunity (Wolter et al., 1993; Peterhansel et al., 1997; Piffanelli et al., 2002; Consonni et al., 2006; Humphry et al., 2006; Kusch et al., 2019). Thus, MLOs have been thought to be host susceptibility factors for PM fungi and act as negative regulators of plant immunity, leading to the inference that *mlo*-mediated resistance is attributable to constitutive or elevated immune responses (Buschges et al., 1997; Acevedo-Garcia et al., 2014). Intriguingly, no genetic components known to function in classical pathogen-associated molecular pattern (PAMP)-triggered immunity (PTI) or effector-triggered immunity (ETI) have been shown to be essential for *mlo*-mediated resistance in Arabidopsis (Consonni et al., 2006; Miklis et al., 2007; Kuhn et al., 2017; Kusch et al., 2019). These observations suggest two possibilities: either MLO proteins repress a potent, yet uncharacterized defense pathway or MLOs are required for a host cellular process critical for PM fungal pathogenesis—such that the observed autoimmunity in *mlo* mutants is a downstream consequence of MLO loss, rather than the primary cause of pathogen resistance.

*MLO* genes in plants belong to a small-to-medium sized gene family with varying members (from a few to a few dozens) that can be divided into seven clades based on protein sequences (Kusch et al., 2016). The genome of *Arabidopsis thaliana* contains 15 *MLO* family members, falling into five clades. Simultaneous impairment of the three clade-V members, *MLO2*, *MLO6* and *MLO12*, results in near-complete resistance to adapted PM pathogens (Consonni et al., 2006). All *MLO* genes whose functional impairment confers PM resistance in dicots belong to clade V, whereas in monocots, such resistance-associated *MLO*s fall within clade IV (Reviewed by Li and Xiao, 2025). The functional equivalence of these two clades is suggested by the absence of clade V *MLO*s in monocots (Kusch et al., 2016). Interestingly, mutations in barley *HvMLO1* (clade IV) or wheat *TaMLO1* (clade IV), or Medicago *MtMLO8* (clade V) cause a reduction in colonization by the arbuscular mycorrhizal fungus *Rhizophagus irregularis* in their respective mutant hosts (Jacott et al., 2020). This suggests that clade IV/V MLOs serve a conserved and important role in accommodating host entry of biotrophic fungi regardless of whether the interaction is mutualistic or pathogenic (Jacott et al., 2021). Several other Arabidopsis *MLO* genes have been shown to be involved in distinct biological processes, including root gravitropism and thigmomorphogenesis (*MLO4* and *MLO11*) (Chen et al., 2009; Bidzinski et al., 2014; Zhu et al., 2021), pollen tube development and guidance (*MLO5*, *MLO9*, *MLO10*, *MLO15*) (Meng et al., 2020; Zhang et al., 2020), and pollen tube reception involving synergid cells (*MLO7*, also known as *NOTIA*) (Kessler et al., 2010; Jones et al., 2017).

All predicted MLO proteins are characterized by an N-terminal region containing seven transmembrane (7-TM) domains and a cytosolic C-terminal tail. This C-terminal region harbors a calmodulin-binding domain (CaMBD), which interacts with calmodulin (CaM) or CaM-like (CML) proteins (Kim et al., 2002; von Bongartz et al., 2023). Several studies have shown that calmodulin binding and/or calcium signaling is required for the full functions of distinct MLOs (Kim et al., 2002; Meng et al., 2020; Zhu et al., 2021; Yuan et al., 2025). Excitingly, several Arabidopsis MLO proteins including MLO2 have been shown to possess calcium channel activity when expressed in animal cells (Cui et al., 2022; Gao et al., 2023), suggesting that MLOs may also play conserved roles in calcium-homeostasis and calcium signaling in plant cells. However, how MLOs’ calcium channel activity is associated with their distinct biological functions in different cells/tissues or subcellular compartments remain unclear.

A GFP-tagged version of barley MLO (HvMLO-GFP) was shown to accumulate focally at the site of cell wall penetration by *Blumeria graminis* f. sp. *hordei* (*Bgh*) in barley epidermal cells (Bhat et al., 2005). This observation led to speculation that HvMLO is actively recruited by the fungus to modulate vesicle-associated processes at the plant cell periphery, thereby facilitating host entry by *Bgh* (Panstruga, 2005). Interestingly, in Arabidopsis, MLO7-GFP is expressed in the synergids of the female gametophyte and is re-localized from Golgi bodies to the filiform apparatus of synergid cells upon arrival of the pollen tube, which is reminiscent of HvMLO’s focal accumulation at the fungal penetration site. Moreover, expressing MLO2-GFP from the *MLO7* native promoter largely restored fertility of the *mlo7* mutant, whereas MLO1-GFP failed to do so (Jones et al., 2017). Interestingly, the chimeric protein NTA-MLO1^CTerm^ (also named faNTA) resulting from swapping the C-terminal CaMBD of MLO1 with that of MLO7 was found to be localized to the filiform apparatus of the synergids, similarly as MLO1, and it fully restored fertility of *mlo7* (Ju et al., 2021). This suggests that the C-terminus of MLO7 specifies its Golgi→ filiform apparatus trafficking upon pollen tube arrival (Ju et al., 2021), which may be regulated through binding of cognate CaM proteins (Yuan et al., 2025). More recently, MLO6 has been shown to interact and colocalize with EXO70H4, a subunit of the exocyst complex, in trichome cells. These two proteins each depends on the other for correct localization and both are required for callose deposition in the trichome cell wall (Huebbers et al., 2024). These findings suggest that MLO6 and likely its two close homologs, MLO2 and MLO12, play an important role in exocytosis of cell wall components (Huebbers et al., 2024).

In this study, we show that loss-of-function mutations in *MLO2*, *MLO6*, and *MLO12* result in a near-complete failure of PM pathogenesis, even in *eds1/pad4/sid2* triple and *eds1/pad4/sid2/pen1/pen2/pen3* sextuple mutant backgrounds, both of which are severely compromised in immune signaling and defense against various (potential) pathogens. These observations suggest that the failure of PM infection in *mlo2/mlo6/mlo12* mutants is not immunity driven. Instead, it likely reflects disruption of a host cellular process essential for PM fungal invasion, which may be co-opted by the pathogen during pathogenesis. We further show that MLO2’s focal accumulation around the penetration peg is determined by its CaMBD-containing C-terminus and that Feronia (FEN) and phosphatidylinositol 4-phosphate 5-kinase 1 (PIP5K1) and PIP5K2 are dispensable for MLO2’s focal accumulation and function that is co-opted by PM fungi for host entry and colonization.

## Results

### Identification of five allelic *MLO2* mutations conferring “resistance” to powdery mildew independent of EDS1, PAD4 and SID2

Plants of the *eps* mutant are super-susceptible to the adapted PM isolate *Golovinomyces cichoracearum (Gc*) UCSC1 (Fig. 1A) (Zhang et al., 2018). To identify host genes crucial for biotrophic pathogenesis of PM fungi, we designed and conducted a large-scale mutant screen in the background of the *eds1/pad4/sid2 (eps)* triple mutant. We aimed to isolate higher-order mutants with compromised immunity yet poor infection (cipi) phenotypes. We reasoned that such *cipi* mutations would more likely disrupt genes essential for PM pathogenesis independent of immunity activation. Eighteen *cipi* mutants were isolated, of which five (*cipi2, cipi3, cipi11, cipi12,* and *cipi15*) showed similar poorest infection by *Gc* UCSC1 (Fig. 1A; Supplementary Fig. S1A). Spore quantification showed that sporulation of *Gc* UCSC1 in *cipi3* plants was reduced to only ∼16% of that in *eps* plants, which is only 1/3 of that in Col-0 (Supplementary Fig. S1B). Interestingly, we noticed that trichomes of those five *cipi* mutants tend to support more fungal growth and sporulation (Fig. 1B). To determine if the causal mutations in the five *cipi* mutants occur in the same gene, we made four crosses between these *cipi* mutants and found that all F_1_ plants showed similar poor infection by *Gc* UCSC1 (Fig. 1C). This result indicates that the causal mutations in the five mutants are allelic. Microscopic examination of PM-infected leaves revealed greatly reduced haustorium formation and conidiophore development, except for occasionally infected trichomes in *cipi3* (Fig. 1D) and other allelic *cipi* mutants. We then crossed *cipi2* with the *eps* parental line and derived an F_2_ segregating population for genetic mapping of the *cipi2* causal mutation. Whole genome sequencing of the bulked segregant pool consisting of 65 F_2_ individuals with *cipi2*-like phenotype revealed a C-to-T synonymous substitution in *MLO2* (At1g11310) in the *cipi2* mutant to be co-segregating with the *cipi* phenotype. This mutation is predicted to create an exonic cryptic donor splice site in the 10th exon of *MLO2* (Fig. 2A), which is predicted to result in a 47 nt deletion of the 10th exon starting from the mutation site, generating a premature stop codon in the 12th exon (Fig. 2A,B). Using the same strategy, a G-to-A synonymous mutation in occurred in *MLO2* of *cipi3* in the acceptor splice site in the 2nd intron (Fig. 2A). This substitution is predicted to activate an immediate downstream exonic cryptic acceptor splice site in the beginning of the 3rd exon, which in turn causes a frameshift resulting in a premature stop codon (Fig. 2C). To check if the transcripts of *MLO2* from these two mutants are indeed mis-spliced, we performed RT-PCR using total RNA extracted from *cipi2* and *cipi3* and found that the transcripts were indeed mis-spliced from *cipi2* or *cipi3* as predicted (Supplementary Fig. S2). The aberrant transcripts are predicted to produce truncated MLO2 proteins of 380 amino acids (*cipi2*) or 97 amino acids (*cipi3*) (Fig, 2D,E). These truncated proteins are probably either nonfunctional or not made due to nonsense-mediated mRNA decay (Baker and Parker, 2004). Next, we sequenced the *MLO2* genomic DNA in *cipi11*, *cipi12* and *cipi15*, and found that each of the three mutants contains a single nonsynonymous exonic mutation in *MLO2*. As depicted in Fig. 2F, *cipi11* contains a G-to-A mutation that changes glycine to arginine (G66R) in the second transmembrane domain, which is identical to the previously reported *mlo2-8* allele (Consonni et al., 2006). *cipi12* contains a G-to-A mutation resulting in a glutamic acid to lysine exchange (E7K) in the N-terminal extracellular domain, which has not been reported before. *cipi15* harbors a G-to-A mutation resulting in an aspartic acid to asparagine substitution (D287N) in the second cytoplasmic loop, which is identical to the *mlo2-11* allele reported before (Consonni et al., 2006).

**Figure 1.**
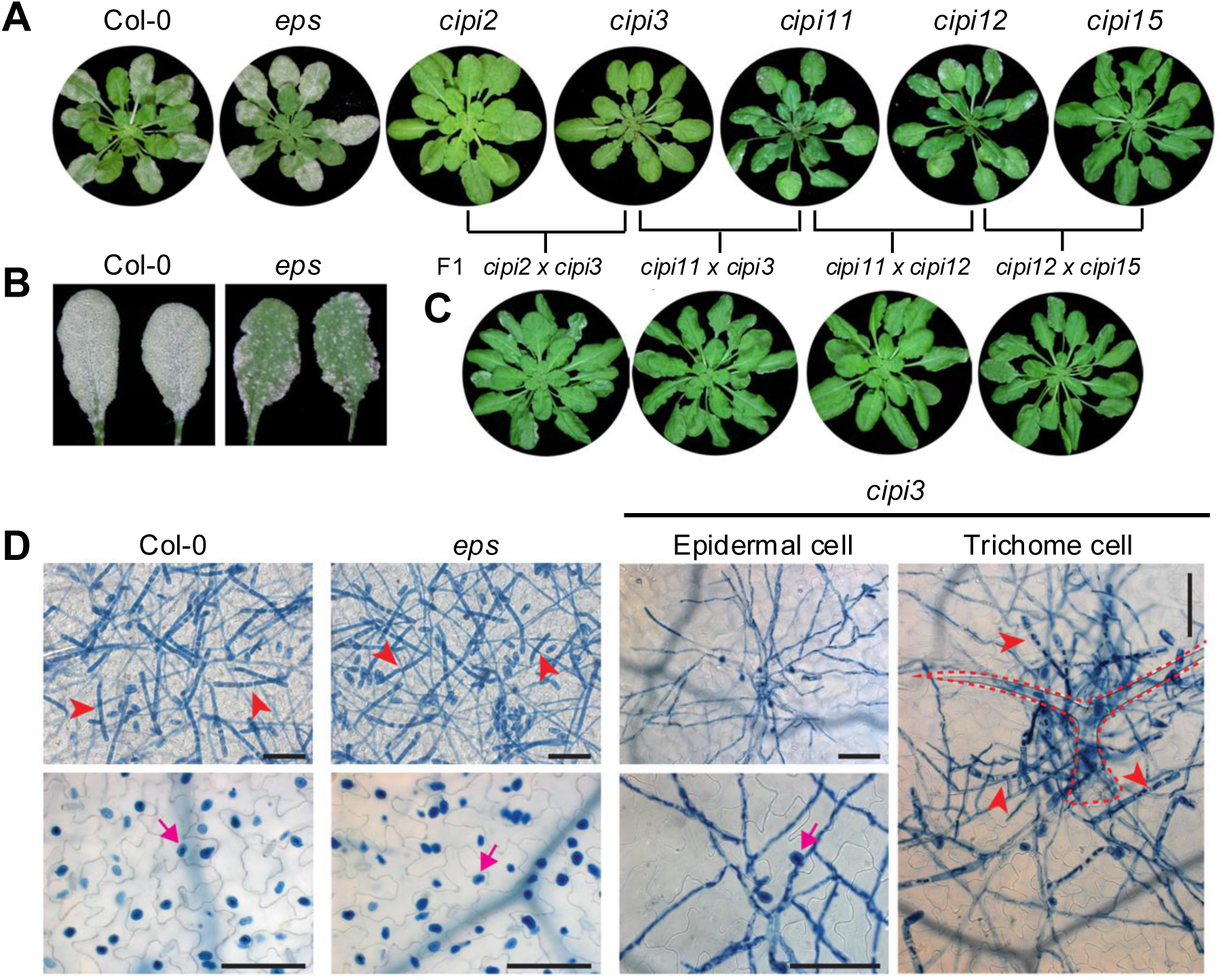
Isolation and phenotypic characterization of five allelic *cipi* mutants. Plants were inoculated with an adapted PM isolate *Golovinomyces cichoracearum* (*Gc*) UCSC1. Photos of infected plants or leaves were taken at 10-15 dpi. **(A)** Representative plants of the five *cipi* mutants along with Col-0 and the *eps* parental line showing their infection phenotypes at 11 dpi. **(B)** Representative leaves from *eps* and *cipi3* at 13 dpi. Note the trichome-based infection in *cipi3*. **(C)** Infection phenotypes of F1 hybrid plants derived from crosses between the indicated mutants at 12 dpi. **(D)** Microscopic images showing fungal structures stained by trypan blue on the leaf surface (upper panel) or inside the epidermal cells (lower panel) of the indicated genotypes at 11 dpi. Note that the trichome cells supporting sporulation in the *cipi3* mutant. Arrowheads indicate rod-shaped conidiophores. Arrows indicate haustoria. Dashed lines highlight a trichome cell. Bar=100 μm.

**Figure 2.**
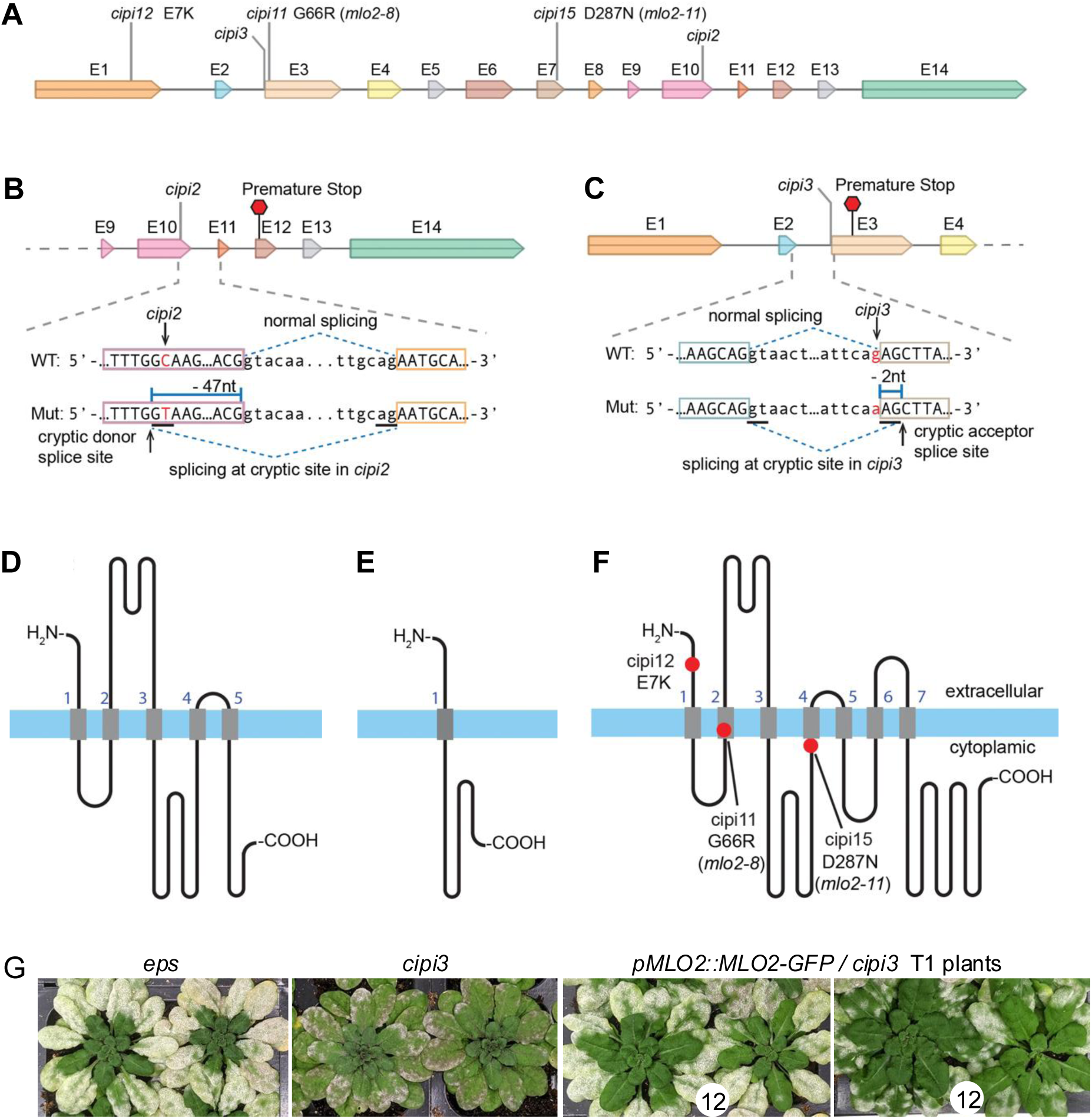
Identification of the five allelic causal mutations in *MLO2*. Bulked segregant pool-based whole genome sequencing identified the candidate causal mutations in *cipi2* and *cipi3* to be in *MLO2*. Targeted sequencing of *MLO2* in *cipi11*, *cipi12* and *cipi15* revealed disruptive mutations as their respective candidate causal mutations. **(A)** Schematic of the *MLO2* gene structure showing the positions of the nucleotide substitutions in the five *cipi* mutants. “E” indicates exon. **(B-C)** Schematic illustration of the positions of the cryptic splice sites (underlined) and the resulting premature stop codons (marked by red hexagons) caused by the mutations in *cipi2* (B) and *cipi3* (C). **(D-F)** Topological illustration of the deduced MLO2 mutant proteins resulting from the *cipi2* (D), *cipi3* (E), and the remaining three *cipi* mutations (F). Highlighted in red dots are three amino acid substitutions in *cipi11*, *cipi12*, and *cipi15* (F). Blue horizontal bars represent membrane bilayers and gray vertical bars represent transmembrane domains. **(G)** Representative plants from the indicated genotypes infected with *Gc* UCSC1 at 12 dpi. Numbers in white circles indicate the number of T1 transgenic plants showing that infection phenotype.

To provide further genetic evidence that mutations in *MLO2* are responsible for the poor infection phenotype of the five *cipi* mutants, we generated transgenic *cipi3* plants expressing *MLO2-GFP* from the native *MLO2* promoter. Half of the 24 T1 *cipi3* plants transgenic for *pMLO2:MLO2-GFP* restored susceptibility to *Gc* UCSC1 to a level close to that of *eps* (Fig. 2H) and the remaining 12 showed medium to low levels of susceptibility. These results further demonstrates that the poor PM infection phenotypes in *cipi3* and other four *cipi* mutants were indeed caused by functional impairment of *MLO2* and that the MLO2-GFP fusion protein is probably fully functional.

Taken together, these genetic data demonstrate that loss of MLO2 significantly reduces PM infection in the *eps* triple mutant background, mirroring the enhanced resistance observed in *mlo2* mutants of Col-0 (Vogel and Somerville, 2000; Consonni et al., 2006). These findings further indicate that the reduced PM infection is independent of EDS1, PAD4, and SID2-mediated immune signaling.

### Loss of *MLO2*, *MLO6* and *MLO12* blocks PM pathogenesis in the *eds1/pad4/sid2* background

The *mlo2-5/mlo6-2/mlo12-1* triple mutants in the Col-0 background (designated *3m/C*) exhibited near complete resistance to PM (Consonni et al., 2006). To see if *3m*-mediated resistance can be recapitulated in the *eps* background, we knocked out *MLO6* and *MLO12* in *cipi3* by CRISPR to create three *eds1/pad4/sid2/mlo2/mlo6/mlo12* (*eps3m*) mutants with indel mutations resulting in early stop codons in *MLO6* and *MLO12* (Supplementary Fig. S3). Infection tests with *Gc* UCSC1 showed that plants of these three sextuple mutants support no visible fungal growth (Fig. 3A). Microscopic examination revealed invariable arrest of PM sporelings shortly after development of the appressorium (arrow in Fig. 3B) and no haustorium formation in the pavement cells (Supplementary Fig. S4). This suggests that the fungus fails to differentiate haustoria from tip of the penetration peg underneath of the appressorium in pavement epidermal cells. Intriguingly, haustorium formation and limited hyphal development could occasionally be found in trichomes (Inset in supplementary Fig. S4), albeit hardly visible to the naked eye, suggesting that the trichome cell environment is different from that of pavement cells. The uninfected or infected plants of the *eps3m* mutants did not show early leaf senescence reported for the *3m/C* triple mutant (Consonni et al., 2006), nor did they exhibited any other obvious developmental phenotypes (Fig. 4A), suggesting that the near-complete resistance to *Gc* UCSC1 is unlikely due to activation of host defense programs.

**Figure 3.**
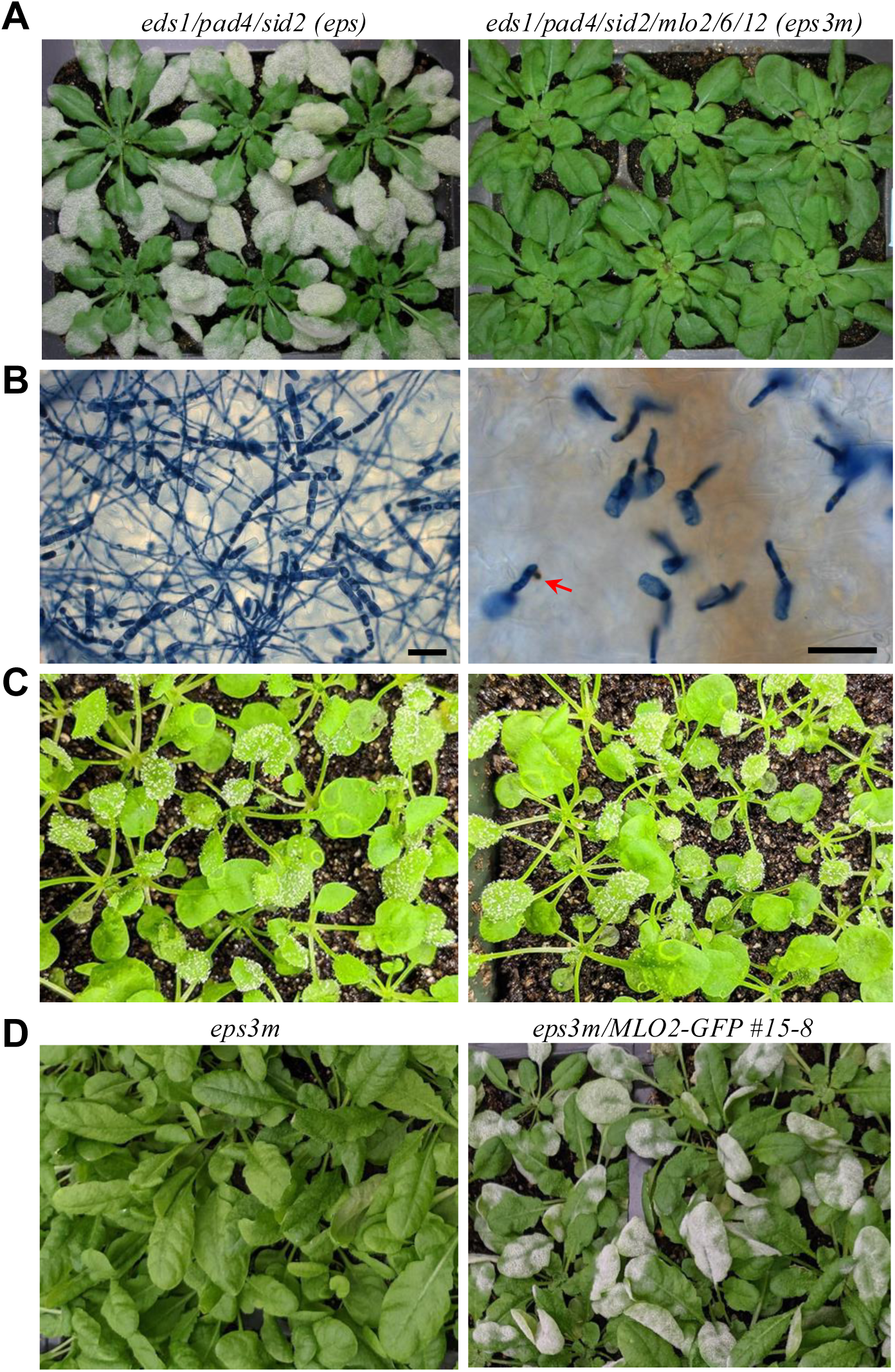
Loss of *MLO2/6/12* in *eps* results in complete lack of infection by powdery mildew. An *eds1-2/pad4-1/sid2-2 mlo2/mlo6/mlo12* sextuple mutant line (*eps3m-7*) was created using CRISPR targeted mutagenesis. Plants of this line were subjected to infection tests and transformation. **(A-B)** Plants of the indicated genotypes inoculated with *Gc* UCSC1. Plant photos (A) or micrographs after trypan blue staining (B) were acquired at 10 dpi or 6 dpi with *Gc* UCSC1, respectively. The arrow indicates penetration peg developed from the appressorium. Bars=50 μm. **(C)** Infection phenotypes of plants of the indicated genotypes inoculated with *a* virulent oomycete isolate *Hyaloperonospora arabidopsidis* Noco2. Photos were taken at 7 dpi. **(D)** *Gc* UCSC1-infection phenotypes of *eps3m* and one *eps3m* transgenic line expressing MLO2-GFP from the *MLO2* promoter. Photos were taken at 10 dpi.

**Figure 4.**
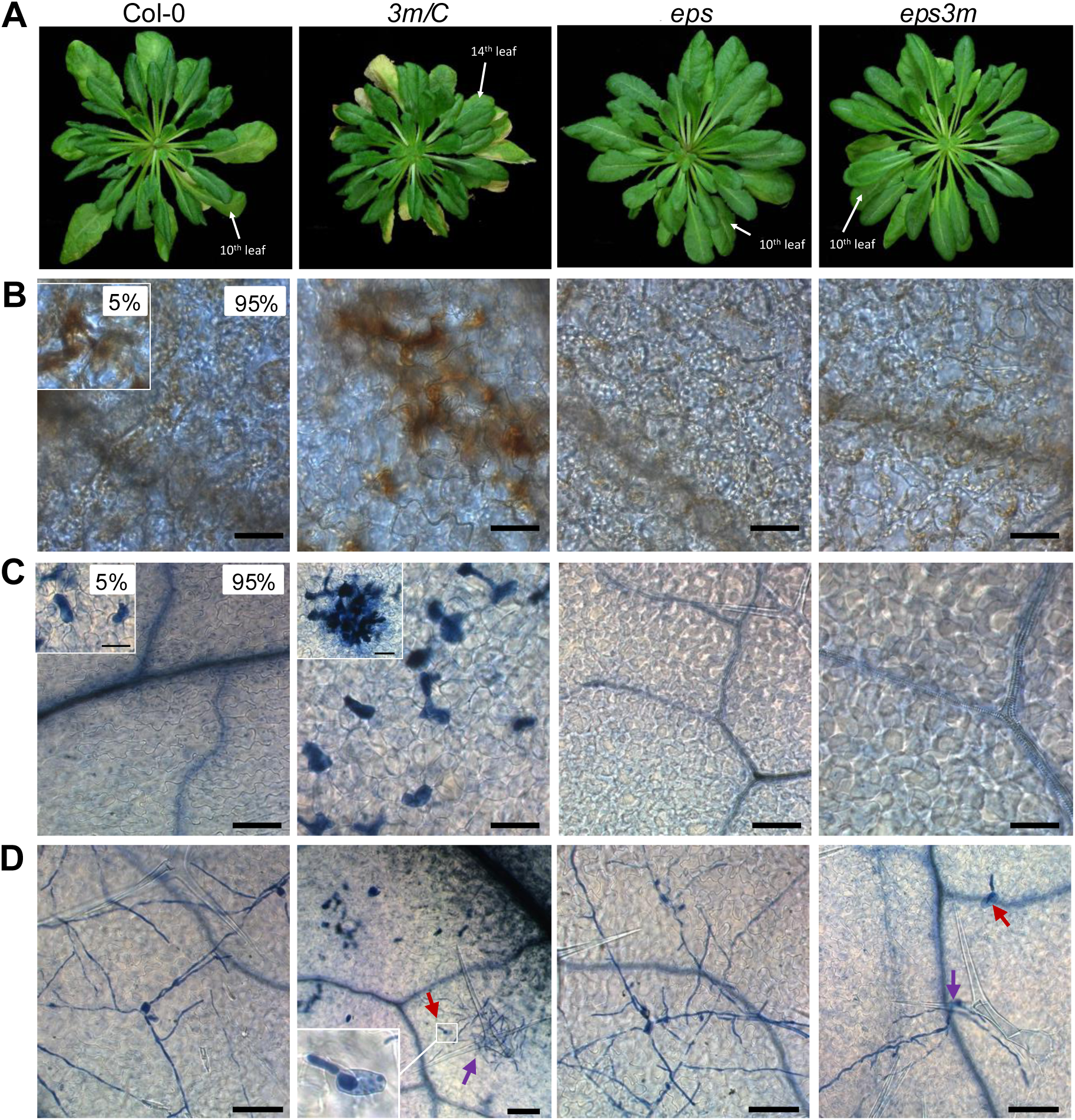
*mlo*-conditioned early leaf senescence is fully suppressed in *eps* and *mlo*-mediated “immunity” is not associated with cell death. Plants of Col-0, *mlo2-5;mlo6-2;mlo12-1* (*3m/*Col-0), *eps* and *eps3m* were grown under short-day for 14 weeks. Selected leaves were subjected for phenotypical analysis. **(A)** Representative plants of the indicated genotypes showing different degrees of leaf senescence (yellowing). **(B-C)** Representative micrographs showing a leaf section of the 10^th^ leaf of Col-0, *eps* and *eps3m*, and the 14^th^ leaf of *3m/*C after 3,3’-diaminobenzidine staining for *in situ* detection of H_2_O_2_ (brownish precipitates in B) or trypan blue staining for cell death (dark blue in C). Bars=50 μm. **(D)** Representative micrographs showing *Gc* UCSC1-infected 11^th^ leaves of Col-0, *eps* and *eps3m*, and the 15^th^ leaves of *3m/*Col-0 plants at 3 dpi after trypan blue staining. Red arrows indicate germinated sporelings that failed to develop further; purple arrows indicate incidental trichome-supported fungal growth. Bars=100 μm.

To further evaluate this inference, we tested *eps* and *eps3m* seedlings with a virulent oomycete pathogen, *Hyaloperonospora arabidopsidis* Noco2. Plants of both genotypes were similarly susceptible to this pathogen (Fig. 3C). This further suggest that the failure of PM fungi in *eps3m* plants is not due to activation of defenses.

We then created *eps3m* transgenic plants expressing *MLO2-GFP* from the *MLO2* promoter and found that such transgenic plants largely restored susceptibility to *Gc* UCSC1 (Fig. 3D), reinforcing the finding that MLO2 is the major contributor to PM susceptibility among the three clade V *MLO* genes (Consonni et al., 2006).

### Failure of PM infection on *eps3m* is not associated with defense activation

To further ascertain whether the near-complete resistance of *eps3m* to *Gc* UCSC1 can be uncoupled from host defense, we first grew plants of Col-0, *3m/C*, *eps*, and *eps3m* under short-day condition for 14 weeks when old rosette leaves of Col-0 wild-type plants started to show weak leaf yellowing (senescence) (Fig. 4A). At this time, plants of *3m/C* showed massive leaf senescence and reduce stature as anticipated, whereas plants of *eps* and *eps3m* showed no sign of leaf senescence (Fig. 4A). Leaf senescence initiation is known to be associated with reactive oxygen species (ROS) production and accumulation in mesophyll cells that eventually leads to collapse of mesophyll and epidermal cells, resulting in leaf yellowing, a typical phenotype of senescence (Jing et al., 2008; Mayta et al., 2019). To see if there is ROS accumulation in leaves of *eps3m* plants, we subjected the 10^th^ leaf from plants of Col-0, *eps* and *eps3m*, and the14^th^ leaf of *3m/C* to 3,3’-diaminobenzidine (DAB) staining for visualization of *in situ* H_2_O_2_ accumulation. While the 10^th^ leaf of the plants of Col-0, *eps* and *eps3m* showed no or little (in the case of Col-0) visible yellowing, the 10^th^ to the 13^th^ leaves of *3m/C* plants exhibited obvious yellowing and their 14^th^ leave showed little yellowing. Subsequent microscopy revealed frequent H_2_O_2_-positve individual and clustered mesophyll cells in the 14^th^ leaves of all six *3m/C* plants examined (Fig. 4B), indicative of the onset of senescence in these leaves. In contrast, such H_2_O_2_-positive cells were completely absent from the 10^th^ leaves of *eps* and *eps3m* plants, and only rarely (∼5% leaf areas) observed in the 10^th^ leaves of Col-0 plants (Fig. 4B), indicating (largely) absence of senescence. We then used trypan blue staining to visualize dead or dying cells in the 10^th^ or 14^th^ leaves of these four genotypes and found very similar patterns as observed for H_2_O_2_ accumulation: while cell death was not detected from the 10^th^ leaves of *eps* and *eps3m* plants, and only rarely seen in the 10^th^ leaves of Col-0 plants, collapse of individual and clustered mesophyll cells, and occasionally more than a dozen of both mesophyll and epidermal cells, was frequently found in the 14^th^ leaves of *3m/C* plants (Fig. 4C).

To examine if PM inoculation can trigger cell death in *3m/C*, and whether the cell death contributes to the termination of fungal development in *3m/C*, we inoculated the detached 11^th^ or 15^th^ leaves of these 14-week-old plants with *Gc* UCSC1 and examined the host-fungal interaction by trypan blue staining at 3 dpi. As shown in Fig. 4D, normal fungal development occurred without cell death in the 11^th^ leaves of the susceptible Col-0 and *eps* plants, whereas the sporelings were completely arrested shortly after germination on pavement cells of both the 15^th^ leaves of *3m/C* and 11^th^ leaves of *eps3m* (indicated by red arrows in Fig. 4D). Notably, while no cell death was observed in *eps3m*, the cell death in *3m/C* was not associated with the early termination of fungal development. As observed before, sporadic hyphal development could be supported by trichome cells in both *3m/C* and *eps3m* (indicated by purple arrows in Fig. 4D).

The above observations suggest that (i) the failure of *Gc* UCSC1 in colonizing host plants due to the loss of *MLO2, MLO6 and MLO12* is independent of ROS production and cell death associated with leaf senescence and (ii) early senescence-associated ROS production and cell death due to the impairment of the three *MLO* genes is *EDS1*, *PAD4* and *SID2*-dependent. The latter agrees with the observation that *mlo2*-conditioned early leaf senescence is *PAD4*- and SA-dependent (Consonni et al., 2006).

Next, we analyzed the expression of four marker genes in mature leaves of seven-week-old short-day-grown plants of Col-0, *3m/C*, *eps*, and *eps3m* prior to and 6, 12 or 48 hrs post-inoculation (hpi) with *Gc* UCSC1. Before inoculation, none of the plants exhibited signs of early senescence. Four marker genes, *FRK1*, *PR1*, *PDF1.2* and *SAG101* were chosen: *FRK1* (Flg22-Induced Receptor-Like Kinase 1) is a marker of PTI activation (Asai et al., 2002), *PR1* induction reports the activation of the SA-dependent defense responses during ETI (and PTI) and systemic acquired resistance (SAR) (Jing et al., 2008; Tsuda et al., 2013), *PDF1.2* is a marker for the activation of the jasmonic acid (JA) and ethylene (ET) pathways (Penninckx et al., 1998), while *SAG101* is a senescence-associated marker gene (He and Gan, 2002). As shown in Supplementary Fig. 5A and B, prior to inoculation with *Gc* UCSC1, expression of *FRK1* in *3m/C* was higher, though not statistically significant, than that in Col-0, suggesting weak constitutive activation of PTI in *3m/C* plants. At 6 hpi, *FRK1* was induced to higher levels in all the four genotypes. At 12 hpi, while the expression of *FRK1* reached significantly higher in *3m/C*, it subsided almost to basal levels in Col-0, *eps* and *eps3m*. The patterns of expression of *PR1* in the four genotypes are similar to those of *FRK1*, except that *PR1* expression in all the four genotypes was low at 6 dpi across the board. These observations suggest that functional impairment of the three *MLO* genes led to pronounced PTI and likely ETI, both of which are mainly EDS1/PAD4/SID2-dependent, as *eps* and *eps3m* had no or little induction of *FRK1* and *PR1* expression before or after inoculation. Expression of *PDF2.1* was barely detectable in plants of all the four genotypes prior to inoculation and induced to higher levels to varied degrees in the four genotypes. However, there was no significant differences among the four genotypes at any time points (Supplementary Fig. S5C). Expression of *SAG101* remained stable and was even slightly reduced after PM inoculation, and there were no significant differences among the four genotypes at any time points (Supplementary Fig. S5D), which is consistent with the lack of any visible leaf senescence in these plants during the period of this experiment.

Taken together, the data shown above reinforce the notion that *MLO2*, *MLO6*, and *MLO12* may negatively regulate *EDS1*/*PAD4*- and SA-associated defense. Importantly, the similar expression of the four marker genes in *eps* and *eps3m* (Supplementary Fig. S5) suggests that *mlo*-mediated ‘resistance’ to PM fungi is unlikely due to the activation of either PTI or ETI.

### MLO2 dosage positively correlates with host susceptibility to *Gc* UCSC1

We noticed that the T1 plants of *eps3m* mutants expressing MLO2-GFP from the native *MLO2* promoter showed varied degrees of susceptibility to *Gc* UCSC1, ranging from near the susceptibility level of *eps* plants to complete lack of infection (Fig. 5A,B). This implies that MLO2 may facilitate PM pathogenesis in a dosage-dependent manner, which could be inferred from the reported differential levels of susceptibility of the transgenic *mlo* lines overexpressing an orthologous wheat or rice *MLO* gene (Piffanelli et al., 2004; Elliott et al., 2005; Ge et al., 2020). To definitively determine if the degree of susceptibility in the T1 lines positively correlates with the levels of MLO2-GFP, we performed a western blot to assess the MLO2-GFP levels using an anti-GFP antibody in *Gc* UCSC1-inoculated plants of the four independent *eps3m-MLO2-GFP* transgenic lines shown in Fig. 5A,B. The full length MLO2-GFP (with an expected size of 93 kDa) was not detectable; however, two smaller bands (∼35 kDa) with ascending intensity were detected in the four lines with increased levels of susceptibility (Fig. 5B-C). Given that GFP is ∼27 kDa, the cleavage products are ∼8 kDa. Because the levels of these two small bands most likely reflect those of the full length MLO2 protein, it is most likely that the MLO2 dosage in those transgenic plants positively correlates with their levels of susceptibility to *Gc* UCSC1. In this context, it is interesting to note that a C-terminal cleavage product was also detected in *E coli-*expressed GST-tagged C-terminus of MLO2 and the cleavage product was diminished when two conserved residues of the C-terminus were mutated (von Bogartz et al., 2023). Whether such cleavage has biological relevance to MLO2’s functionality remains to be determined.

**Figure 5.**
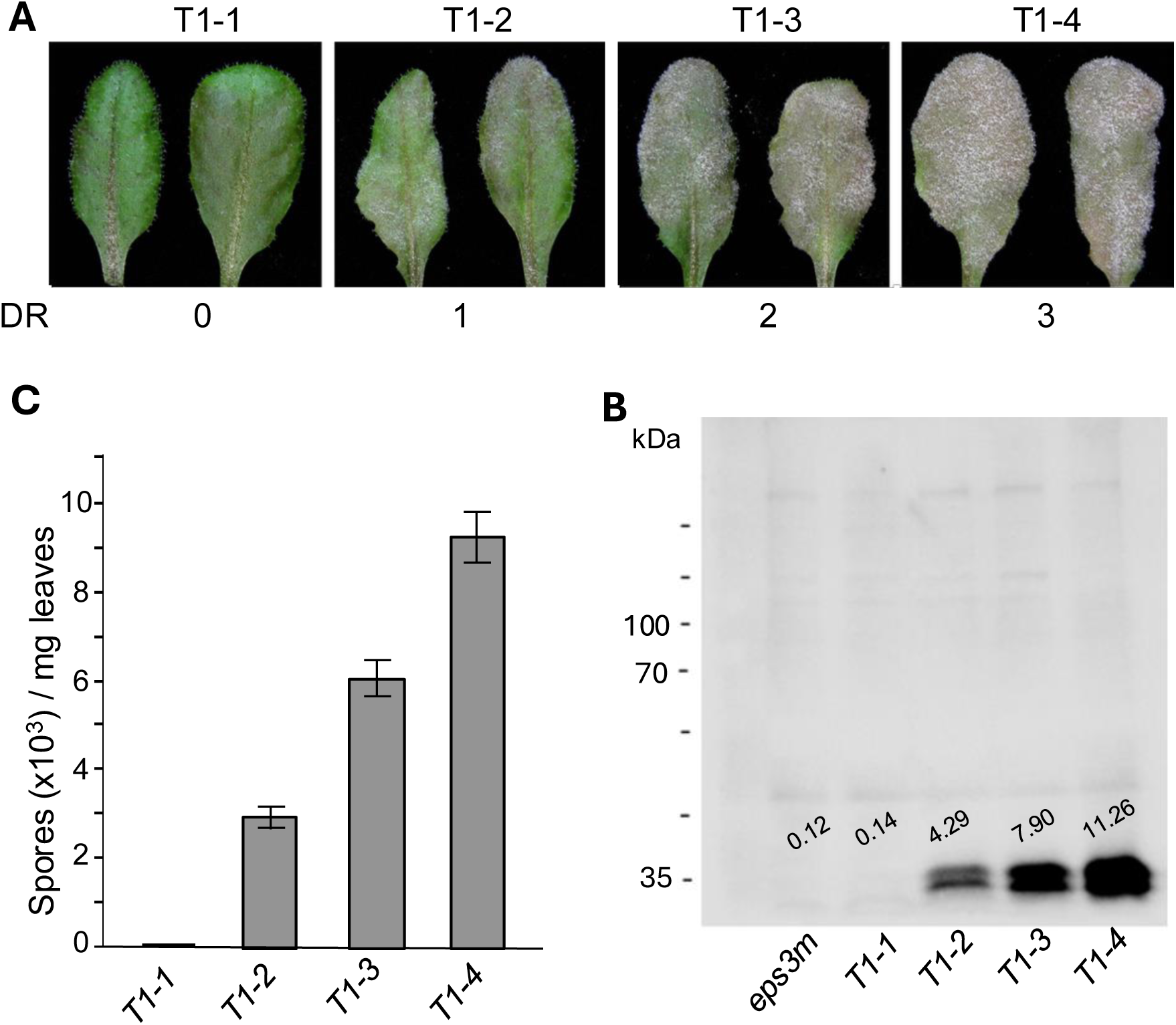
MLO2-GFP expression levels positively correlate with susceptibility to *Gc* UCSC1 **(A)** Representative leaves of four selected *epsm3* lines transgenic for *MLO2p::MLO2-GFP* showing different levels of susceptibility to *Gc* UCSC1. Photos were taken at 11 dpi. DR, disease reaction score judged visually based on leaf coverage of fungal mass. **(B)** Quantification of total number of spores per mg of infected leaf tissues of the four lines. Different letters indicate statistically significant differences (*P*<0.05) between the three lines, as determined by multiple comparisons using one-way ANOVA, followed by Tukey’s HSD test. This experiment was repeated once with similar results. **(C)** Western blot using anti-GFP antibody to measure MLO2-GFP protein levels in the four lines. The *eps3m* parental line served as negative control. Numbers above the bands indicate intensity of the two bands against the blank background measured with ImageJ.

### ER/Golgi-localized MLO2 is targeted to peri-penetration peg membranous space

Barley HvMLO-GFP and Arabidopsis MLO2-GFP have been shown to accumulate at the fungal penetration site (Bhat et al., 2005; Qin et al., 2020). Because the term “penetration site” is rather vague, we sought to determine MLO2’s subcellular localization with higher spatiotemporal resolution to better understand the cellular functions of MLO2 in facilitating PM pathogenesis. To this end, we first inoculated plants of the *eps3m*-*MLO2-GFP* transgenic line #4 with *Gc* UCSC1 for localization analysis. A close microscopic examination of infected leaves stained with propidium iodide at 2 dpi revealed accumulation of MLO2-GFP in a collar-like structure around the penetration peg underneath the appressorium, with MLO2-GFP puncta distributing around the penetration site (Fig. 6A, B). To better examine MLO2-GFP’s dynamic spatiotemporal localization during appressorium-haustorium differentiation, we introduced the *pRPW8.2::RPW8.2-RFP* construct into the *eps3m*-*MLO2-GFP* background. RPW8.2 is specifically targeted to the extra-haustorial membrane (EHM) encasing the haustorium that emanates from the tip of the penetration peg, hence RPW8.2 can serve as a reporter of the spatiotemporal biogenesis of the EHM and the haustorium (Wang et al., 2009). A time course analysis showed that PM spores germinated at ∼6 hpi and MLO2-GFP first exhibited detectable focal accumulation at 7.5 hpi (Fig. 6C,D) whereas RPW8.2-RFP was first detectable at the EHM around 16 hpi (Fig. 6E) (Wang et al., 2009). Importantly, MLO2-GFP was observed to accumulate in a compartment (∼1-2 μm thick and 1-3 μm long) surrounding the penetration peg. We tentatively named this compartment the <u>p</u>eri-<u>p</u>enetration-peg <u>m</u>embranous space (PPM). Notably, MLO2-GFP was never found in the EHM which initiates from the haustorial neck connecting to the penetration peg (Fig. 6F-H; Supplementary Fig. S6). These observations imply that MLO2 is specifically recruited to the <u>p</u>eri-<u>p</u>enetration-peg <u>m</u>embranous space (PPM) (Fig. 6I,J) where it may play an important role in sealing and stabilizing the convoluted membrane junction, thereby accommodating haustorium differentiation (See Discussion).

**Figure 6.**
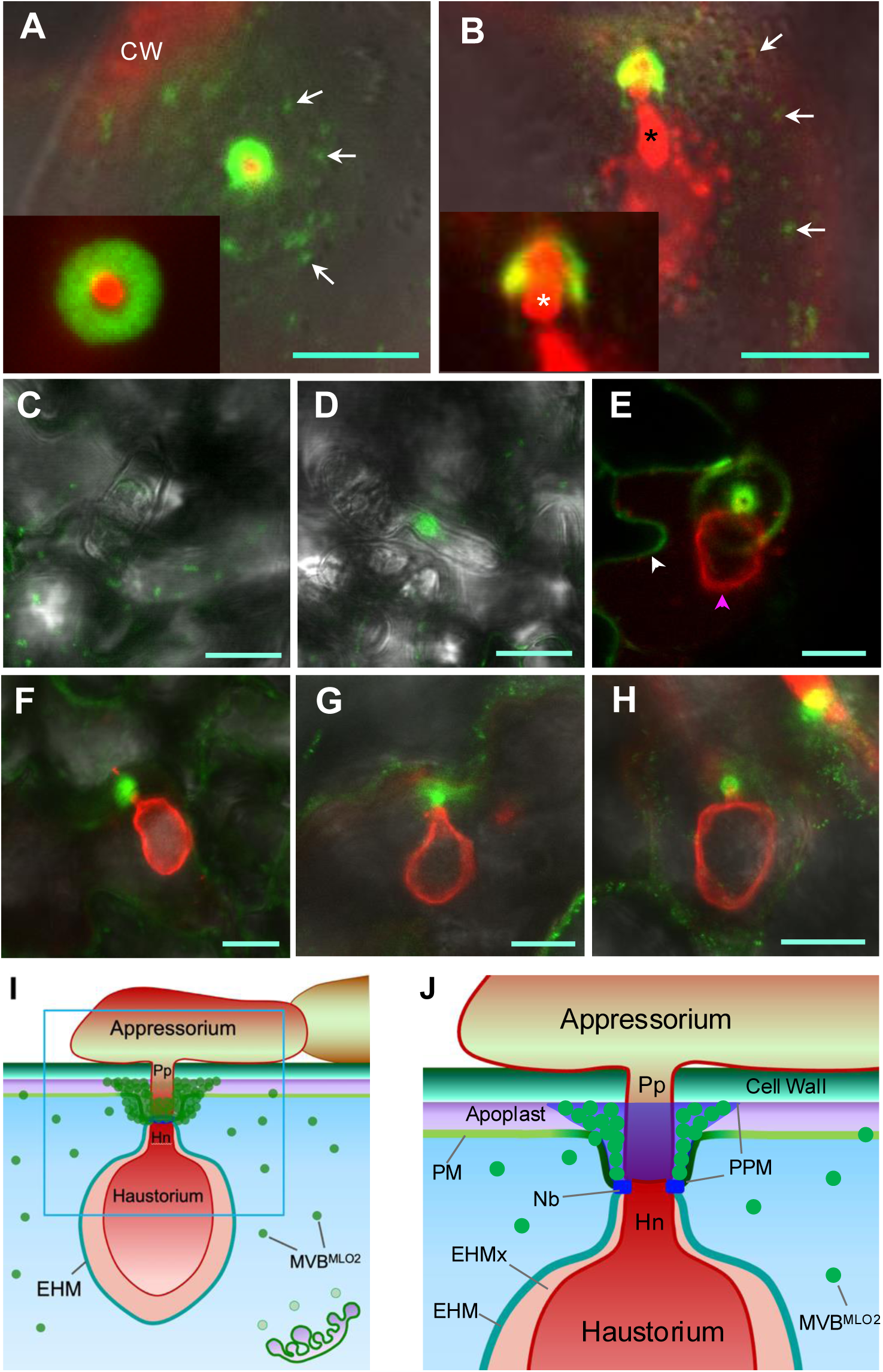
MLO2-GFP is localized to the peri-penetration peg membranous space (PPM). Plants of *eps3m* expressing MLO2-GFP (A,B) or plants of *eps3m* expressing both MLO2-GFP and RPW8.2-RFP (C-H) were used to determine MLO2-GFP’s localization by confocal imaging. All images are Z-stack projections of 5 to 15 optical sections. Bar=10 μm. **(A,B)** Representative images showing localization of MLO2-GFP to the plasma membrane of a leaf epidermal cell penetrated by *Gc* UCSC1. Image in (A) is a top-down view of a fungal penetration site, whereas image in (B) is a side view of the same site. Insets are closeup view of a single optical section. The fungal structure is stained red by propidium iodide. White arrows indicate MLO2-GFP puncta which may be MVBs or endosomes. White asterisk indicates penetration peg, while dark asterisk indicates haustorial neck. CW, cell wall. **(C)** A representative image showing punctum distribution of MLO2-GFP at 6 hpi. **(D)** A representative image showing focal accumulation of MLO2-GFP at 7.5 hpi. **(E)** A representative image showing localization of MLO2-GFP and RPW8.2-RFP at 16 hpi. White arrowhead indicates the plasma membrane, while magenta arrowhead indicates the extra-haustorial membrane (EHM). **(F-H)** Images showing MLO2-GFP accumulates around the penetration peg, next to the haustorial neck (also see Supplementary Fig. S4). **(I)** A cartoon depicting the focal accumulation of MLO2 from Golgi bodies to the penetration site. Pp, penetration peg; Hn, haustorial neck; MVB^MLO2^, multivesicular bodies (MVBs) containing MLO2; EHM, extra-haustorial membrane. **(J)** A zoom-in cross section of (I) illustrating the membrane junction where MLO2 focally accumulates. We propose that exosomes derived from MVBs^MLO2^ are secreted into the space between the penetration peg and the host cell wall and plasma membrane (PM) to seal this membrane junction. We designate this junction filled with MLO2-containing exosomes or paramural bodies as the <u>p</u>eri-<u>p</u>enetration peg <u>m</u>embranous space (PPM). PPM may also include the perturbed PM and the haustorial neckband (Nb), which is believed to be a diffusion barrier between the apoplast and the extra-haustorial membrane matrix.

### Ectopic expression of *MLO7* but not *MLO1* partially restores susceptibility of *eps3m* to *Gc* UCSC1

Several *MLO* family members in Arabidopsis exhibit distinct expression patterns and serve distinct biological functions (Chen et al., 2006; Davis et al., 2017; Jones and Kessler, 2017; Li and Xiao, 2025). Given that MLO1, MLO2 and several other MLOs assayed possess calcium channels activity when ectopically expressed in mammalian cells (Gao et al., 2022; Gao et al., 2023), we wondered if MLOs belonging to other clades and expressing in other organs/trissues share the same molecular functions as MLO2 in facilitating PM pathogenesis when ectopically expressed in leaf epidermal cells. Since the *eps* triple mutant is super-susceptible to *Gc* UCSC1, whlie *eps3m* is essentially “immmue” to *Gc* UCSC1, we reasoned that any functional complementation of the loss of *MLO2/6/12* in *eps3m* should be readily manifested by fungal growth visible to the nake eye. To test this, we made *eps3m* transgenic plants expressing barley *MLO1* (*HvMLO1*) from the *35S* promoter and found that nine of 20 T1 transgenic plants supported varied degree of susceptibility to *Gc* UCSC1 (Supplementary Fig. S7A), indicating that HvMLO1 can largely perform the same molecular function as MLO2 in Arabidopsis, despite that it belongs to a different clade (clade IV) in the MLO family tree (Kusch et al., 2016). We then made transgenic *eps3m* lines expressing MLO6-GFP from the *MLO6* promoter. Five of 14 *Gc* UCSC1-infected T1 plants transgenic for *MLO6-GFP* showed visible but limited fungal growth (Fig. 7A) with ∼10% of spore-production compared to those expressing *MLO2-GFP* from the *MLO2* promoter (Supplementary Fig. S7B,C). Confocal imaging showed that MLO6-GFP also exhibited similar focal accumulation at the fungal penetration site in the five transgenic lines (Fig. 7B). These results support the notion that *MLO2* play a major role while *MLO*6 and *MLO12* play a minor role in permitting PM pathogenesis (Consonni et al., 2006) and that protein accumulation at PPM may be a common feature of MLOs that function as susceptibility factors. The above results also demonstrate that PM infection in *epsm3* transformants can be used as a sensitive reporter to evaluate whether any candidate wild-type or mutant *MLO* genes can perform the same molecular function as *MLO2* by ectopically expressing it in leaf epidermal cells from the *MLO2* promoter.

**Figure 7.**
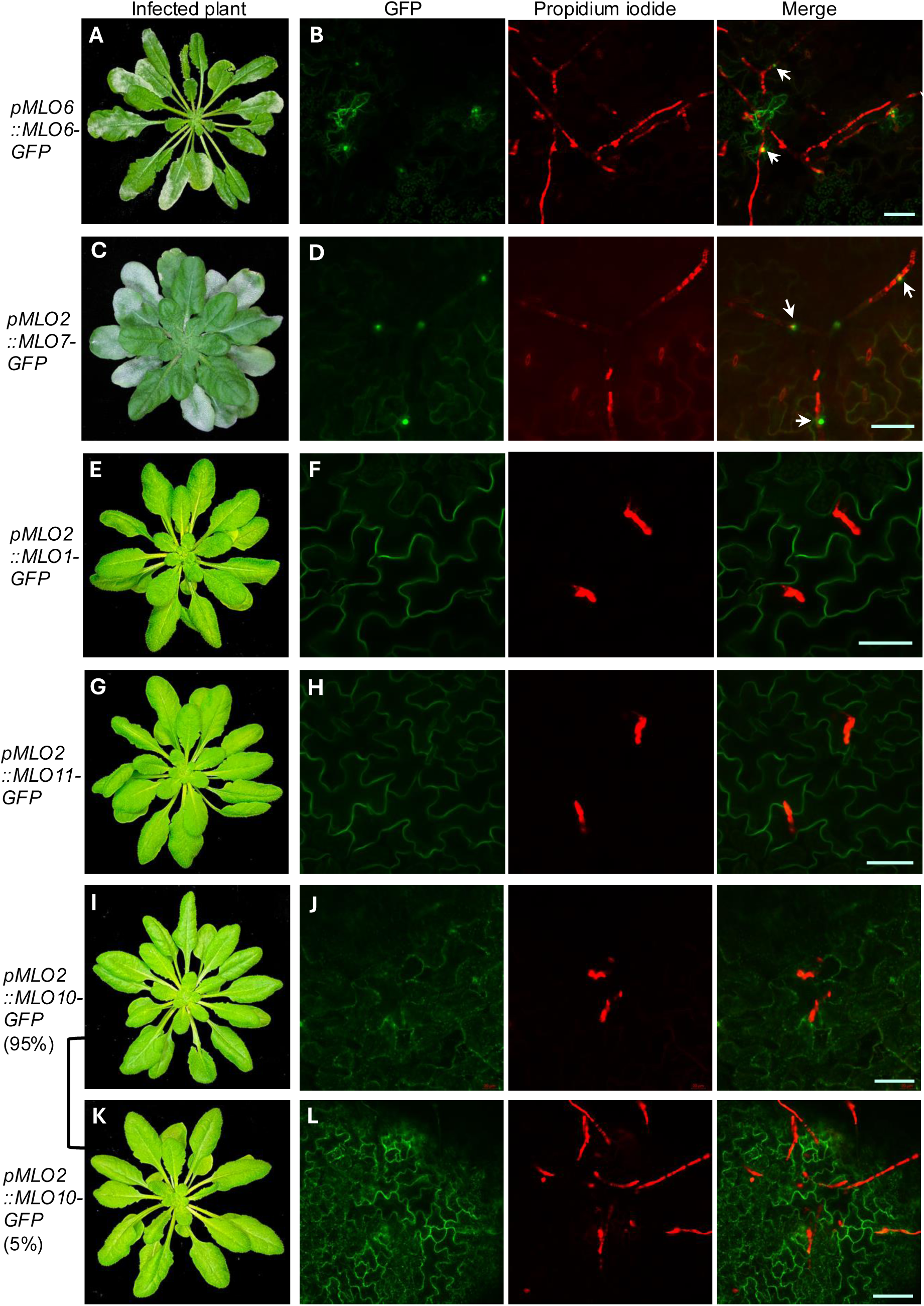
Functional complementation tests for other MLO family members in *eps3m*. Plants of *eps3m* lines transgenic for the indicated *MLOx-GFP* DNA constructs were inoculated with *Gc* UCSC1. Infection phenotypes were visually and microscopically examined. Plant photos were taken at 10-11 dpi. Infected leaves were subjected to confocal imaging at 3 dpi for assessing the localization of each MLOx-GFP fusion protein. Images shown are Z-stack projection of 3-5 optical sections. Fungal structures were stained red with propidium iodide. Arrows indicate penetration sties. Bar=50μm.

A similar strategy was employed to show that expression of *MLO2* in synergid cells from the *MLO7* (*NTA*) promoter significantly restored the fertility of the *mlo7* mutant (Jones et al., 2017), suggesting that MLO2 and MLO7 share similar molecular function. To provide concrete evidence for the converse scenario, we generated *eps3m* plants expressing MLO7-GFP from the *MLO2* promoter and found that five of 18 T1 plants were moderately susceptible and six T1 plants were weakly susceptible to *Gc* UCSC1 (Fig., 7C; Supplementary Fig., S7D). Not surprisingly, like MLO2-GFP, MLO7-GFP also exhibited focal accumulation at the PPM (Fig. 7D). This indicates that MLO7 can partially complement the loss of *MLO2/6/12* if ectopically expressed in leaf epidermal cells.

To further expand such functional assays to other MLO clades, *MLO4* and *MLO11* (belonging to clade I), *MLO1* (belonging to clade II), *MLO5* and *MLO10* (belonging to clade III), and *MLO3* (belonging to clade VI) were cloned into the same binary vector that contains the *MLO2* promoter for translational fusion with *GFP* (Supplementary Fig. S8). At least eight *eps3m* T1 transgenic plants were produced for each construct. All T1 plants were inoculated with *Gc* UCSC1 to determine if (i) any plants can support fungal growth and sporulation and (ii) whether the fusion proteins are detectable by confocal microscopy and if so, where they are localized.

Intriguingly, none of the T1 plants expressing any of the six tested *MLO* genes (*MLO1*, *MLO3*, *MLO4*, *MLO5*, *MLO10* and *MLO11*) supported visible growth of *Gc* UCSC1 (Fig. 7E, G, I, K) and no GFP signal was reliably detected in the T1 plants transgenic for *MLO3-GFP*, *MLO4-GFP* and *MLO5-GFP* (*not shown*). Among the 60 *eps3m* T1 plants expressing *MLO1-GFP*, GFP signal was readily detected in the plasma membrane (labeled by the lipophilic dye FM4-64) of leaf epidermal cells (Supplementary Fig. S9A), which is consistent with the MLO1-GFP localization pattern observed in epidermal cells of *Nicotiana benthamiana* leaves after agrobacterium-mediated transient expression (Jones and Kessler, 2017). Notably, sporelings were arrested probably due to failure in host entry in *eps3m* plants expressing MLO1-GFP and there was no detectable change of MLO1-GFP in response to attempted fungal penetration (Fig. 7F). Similarly, GFP signal was detected in the plasma membrane of the leaf epidermal cells in the eight *eps3m* T1 plants transgenic for MLO11-GFP (Supplementary Fig. S9B) and the sporelings failed to develop further (Fig. 7H). Interestingly, although no fungal mass was visible to the naked eye in any of the 15 *eps3m* T1 plants transgenic for *MLO10-GFP* (Fig. 7I and K), a few small fungal colonies with very limited hyphal growth without conidiophore formation were detectable in T2 progenies of two independent T1 lines (Fig. 7L). MLO10-GFP was detected in puncta and likely at the plasma membrane, but there was no obvious focal accumulation even in the case of successful penetration reported by limited mycelial growth (Fig. 7L). These observations suggest that in term of subcellular localization, MLO10 is similar to MLO2 but distinct from MLO1 and MLO11. Given that MLO10 was able to restore fertility of *mlo7* when expressed in synergid cells (Jones et al., 2017), the inability of MLO10-GFP to significantly complement PM susceptibility in the *eps3m* background was unexpected, especially given its close homology to MLO7.

To validate the three MLO-GFP fusion constructs, i.e., *MLO3-GFP*, *MLO4-GFP* and *MLO5-GFP,* whose expression was not detectable in their respective Arabidopsis transgenic plants, and assess their subcellular localization, we transiently co-expressed them with HDEL-mCherry (an ER marker) and Man1 (1-49)-mCherry (a Golgi marker) (Nelson et al., 2007) in *N. benthamiana* leaves via agroinfiltration. Confocal imaging at three days post-infiltration revealed that all three fusion proteins were detectable and exhibited similar patterns of partial co-localization with HDEL-mCherry and/or Man1-mCherry (Supplementary Fig. S10–12). These observations suggest that MLO3-GFP, MLO4-GFP, and MLO5-GFP are also likely localized to the ER and/or Golgi compartments in *Arabidopsis*; however, their low expression and/or rapid turnover may prevent reliable detection in stable *eps3m* transgenic plants.

### Domain-swapping between MLO1 and MLO2 reveals the C-terminal CaM-binding domain as a key determinant of subcellular localization

MLO7 (i.e., NTA) localizes to Golgi bodies in synergid cells and redistributes to the plasma membrane at the filiform apparatus during pollen tube reception (Jones et al., 2017; Ju et al., 2021). A chimeric MLO protein that contains the N-terminal 7TM portion of NTA and the cytoplasmic CaMBD-containing C-terminus of MLO1 (designated faNTA) exhibits constitutive localization at the filiform apparatus and is able to restore the fertility of *nta* (Ju et al., 2021). This finding suggests that the C-termini of MLO1 and MLO7 determines their respective localization. To determine whether the CaMBD-containing C-termini of MLO1 and MLO2 dictate their respective subcellular localization in leaf epidermal cells, we constructed two chimeric genes by swapping the fragments encoding their C-terminal domains. The first resulting chimeric protein consists of MLO1’s N-terminal 7TM (1-438 amino acids) and MLO2’s C-terminus (442-574 amino acids) (designated MLO1n-2c), while the second comprises MLO2’s N-terminal 7TM (1-442 amino acids) with MLO1’s C-terminus (439-526) (designated MLO2n-1c) (Fig. 8A). These chimeric genes, *MLO1n-2c* and *MLO2n-1c*, were translationally fused with GFP at their C-termini, and the resulting fusion constructs were stably expressed in *eps3m* from the *MLO2* promoter. More than 20 *eps3m* T1 plants transgenic for either of the two chimeric fusion genes were generated and inoculated with *Gc* UCSC1. None of the 23 T1 plants transgenic for *MLO1n-2c-GFP* supported any visible fungal growth and propidium iodide staining showed that sporelings were arrested shortly after germination (Fig. 8B), and MLO1n-2c-GFP exhibited punctum distribution (Fig. 8C). To test if successful penetration followed by haustorial differentiation can induce MLO1n-2c-GFP’s focal accumulation, we introduced the same *MLO2p::MLO1n-2c-GFP* construct into the *eps* background to allow growth of *Gc* UCSC1. The T1 transgenic plants were as susceptible as *eps* (Fig. 8D), suggesting that MLO1n-2c expression has no dominant negative effect. Interestingly, MLO1n-2c-GFP was found to accumulate at the PPM in the *eps* transgenic plants (Fig. 8E). By contrast, seven out of 21 e*psm3* T1 plants transgenic for *MLO2n-1c-GFP* supported fungal hyphal growth as revealed by propidium iodide staining at 3 dpi (Fig. 8G), and three of them also supported weak *Gc* UCSC1 infection visible to the naked eye at 13 dpi (Fig. 8F). Interestingly, similar to MLO1-GFP (Fig. 8F), MLO2n-1c-GFP exhibited plasma membrane localization in leaf epidermal cells, and there was no obvious focal accumulation around the fungal penetration site (Fig. 8G). Collectively, the above results indicate that the C-terminus of MLO1 confers plasma membrane localization whereas the C-terminus of MLO2 endows punctum distribution and fungal penetration-induced focal accumulation at the PPM. The results also suggest that (i) the N-terminal 7TM portion of MLO2, but not that of MLO1, specifies the function of MLO2 in accommodating entry of the PM pathogen and (ii) the full function of MLO2 requires its enrichment at the PPM, which is regulated by its CaMBD-containing C-terminus.

**Figure 8.**
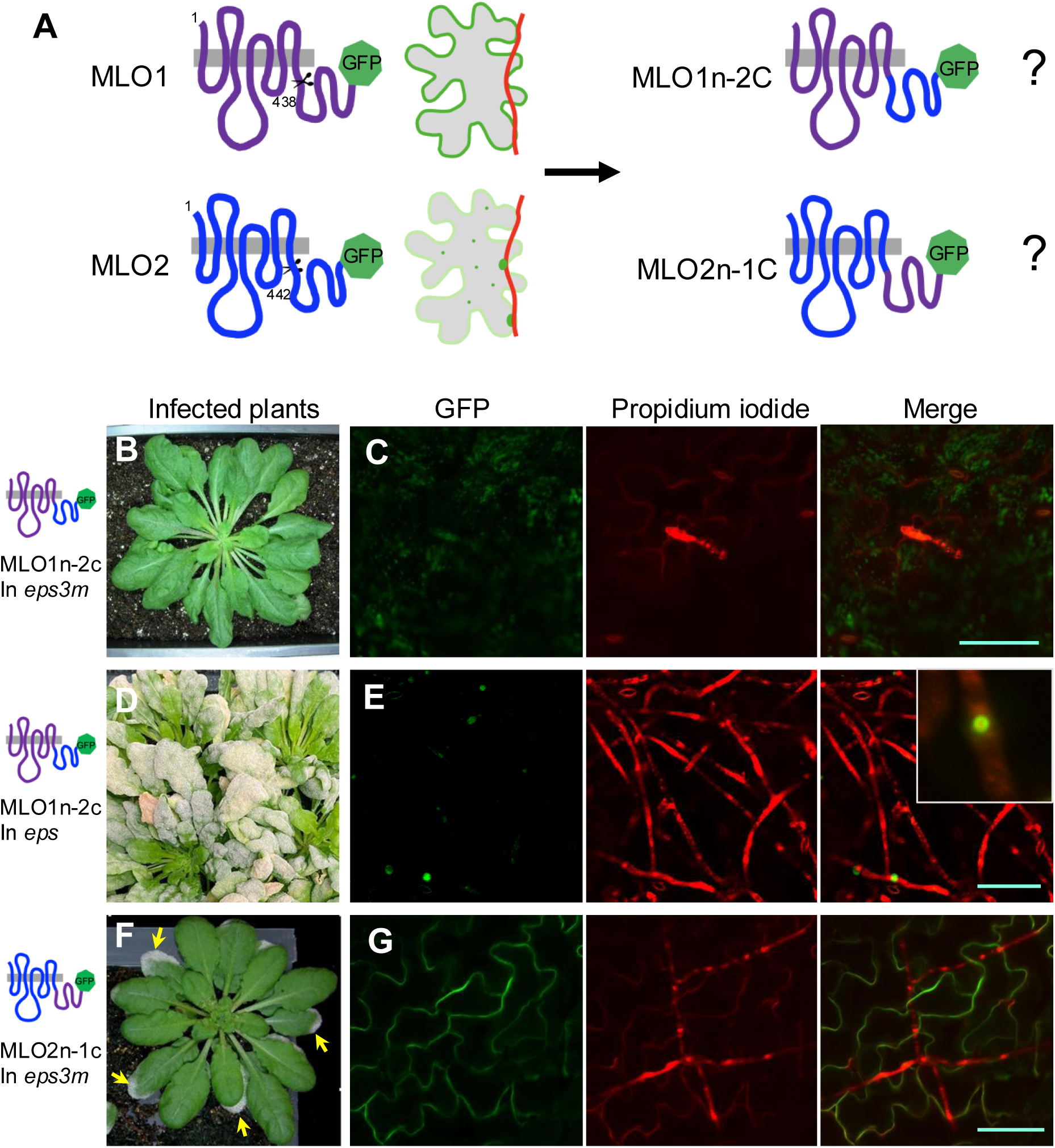
Domain-swapping analysis between MLO1 and MLO2 reveals two functional arms of MLO2. **(A)** Schematic illustration of the experimental design. **(B,D,F)** Infection phenotypes of representative plants expressing MLO1n-2c-GFP or MLO2n-1c-GFP from the *MLO2* promoter in either the *eps3m* background (B,F) or the *eps* background (D). Photos were taken at 12 dpi with *Gc* UCSC1. Arrows in (F) point to infected leaf tips with visible fungal mass. **(C,E,G)** Representative confocal images showing localization of the indicated chimeric fusion proteins in leaf epidermal cells at 3 dpi. Images are Z-stack projection of 3-5 optical sections. Fungal structures were stained red with propidium iodide. Inset in E is a closeup view of a single penetration site where MLO1n-2c accumulates. Bar=50μm.

### Loss of *FERONIA* does not affect MLO2’s localization and role in facilitating PM pathogenesis

FERONIA (FER), a member of the *Catharanthus roseus* receptor-like kinase 1-like (CrRLK1L) protein subfamily (Escobar-Restrepo et al., 2007; Guo et al., 2009), has been shown to be important for the proper localization of MLO7 in synergid cells (Ju et al., 2021). Additionally, *fer* mutants exhibit reduced susceptibility to PM disease (Kessler et al., 2010). Given the conserved molecular functions shared by MLO2 and MLO7 (Fig. 7C; Jones et al., 2017), we asked whether FER is also required for the focal localization of MLO2 at the PPM in leaf epidermal cells. To test this, we first performed CRIPSR-targeted mutagenesis in the *eps* background to determine if *FER* is required for MLO2 function and if so, *eps-fer* mutants should exhibit reduced susceptibility to *Gc* UCSC1 as seen in the five *mlo2 cipi* mutants (Fig. 1). Loss-of-function *fer* mutants displays a more compact rosette phenotype readily distinguishable from wild-type plants (Deslauriers and Larsen, 2010). We obtained seven presumable *eps*-*fer* mutants with a compact rosette and found that all of them showed similar levels of susceptibility to *Gc* UCSC1 as *eps* plants (Fig. 9A). Sequencing three of the seven mutants revealed disruptive indel mutations close to the sgRNA target sites in the second exon of *FER* (Supplementary Fig. S13A). This result indicates that, unlike *MLO2*, *FER* is dispensable for PM pathogenesis in *eps* and implies that the reported enhanced resistance in *fer* single mutants (Kessler et al., 2010) may be due to activation of EDS1/PAD4/SID-dependent immunity. Next, we introduced the same CRISPR/Cas9 construct into *eps3m* plants transgenic for *MLO2-GFP* and *RPW8.2-RFP* to determine if focal accumulation of MLO2-GFP is affected in the absence of FER. Among 27 T1 transgenic plants obtained, 18 displayed the compact rosette phenotype. Among the 18 plants, eight were susceptible to *Gc* UCSC1, the remaining10 plants did not support any growth of *Gc* UCSC1 visible to the naked eye (Fig. 9B). Targeted sequencing of *FER* in three of the eight susceptible plants with a compact rosette identified disruptive indels in *FER* as expected (Supplementary Fig. S13B). Confocal imaging showed that MLO2-GFP signal was not detectable in the 10 T1 plants with a compact rosette but lacking infection, suggesting that the *MLO2-GFP* transgene is silenced, which was frequently observed in transgenic plants generated in the *eps3m*-*MLO2-GFP;RPW8.2-RFP* background (data not shown). Importantly, MLO2-GFP, as well as RPW8.2-RFP in the eight T1 plants susceptible to the fungus exhibited normal localization (Fig. 9C,D), indicating that loss of FER does not affect MLO2’s localization to the PPM and its role in accommodating PM pathogenesis in leaf epidermal cells. To assess if there is functional redundancy among FER and its family members, we performed multiplexed CRISPR to target eight *CrRLK1L* family members including *FER* that are known to be involved in immunity and/or expressed in leaves based on the search results from the RNA-seq database which include >20,000 RNA-seq libraries (https://plantrnadb.com/athrdb/). Among 25 T1 transgenic lines, 11 displayed the compact rosette phenotype, among which five were susceptible (Fig. 9E). We selected one of the susceptible lines for sequencing of all the eight targeted *CrRLK1L* genes. The results showed that this line contains disruptive mutations in *FER* and five additional *CrRLK1Ls* (Fig. 9F; Supplementary Fig. S14), yet MLO2-GFP’s localization and role in accommodating PM entry remain unaffected (not shown), indicating that those six CrRLK1Ls are dispensable for MLO2’s localization and function.

**Figure 9.**
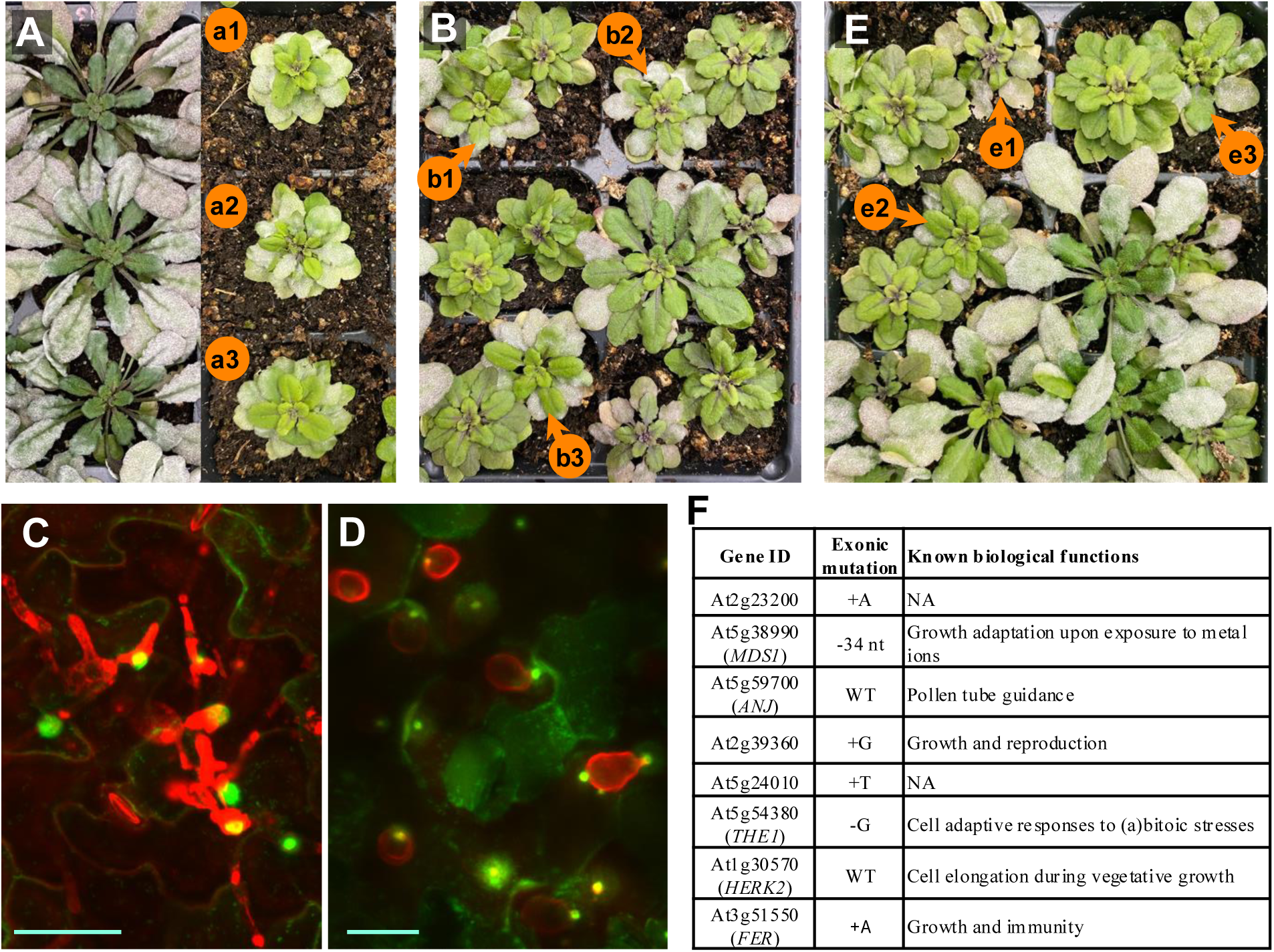
FERONIA and its five family members are dispensable for MLO2’s focal accumulation and MLO2-mediated susceptibility. CRISPR-targeted mutagenesis was used to knock out *Feronia* (*FER*) alone, or *FER* and other five members of the *CrRLK1L* gene family in *eps* or *eps3m* expressing MLO2-GFP and RPW8.2-RFP. Independent T1 plants were inoculated with *Gc* UCSC1 and plant photos were taken at 10-12 dpi. Mutations in *FER* and other family member were identified by targeted sequencing (see Supplementary Fig. S10 and S11). **(A)** Infection phenotypes of representative *eps* plants and three T1 transgenic lines of *eps3* expressing the CRISPR construct targeting *FER* (a1-a3). Sanger sequencing results are shown in Supplementary Fig. S10A) **(B)** Infection phenotypes of T1 plants of *eps3m/pMLO2-MLO2-GFP/pRPW8.2-RPW8.2-RFP* transgenic for the CRISPR construct targeting *FER*. Three susceptible *fer-*like mutant plants (b1-b3) were subjected to sequencing analysis of *FER* (see Supplementary Fig. S10B). **(C,D)** Representative confocal images showing typical focal accumulation of MLO2-GFP (C) and EHM-localization of RPW8.2-RFP (D). Images are Z-stack projection of 3-5 optical sections. Fungal structures were stained red with propidium iodide. Bar=20μm. **(E)** Infection phenotypes of T1 plants of *eps3m/pMLO2-MLO2-GFP/pRPW8.2-RPW8.2-RFP* transgenic for the CRISPR construct targeting eight *CrRLK1L* family genes. Three susceptible *fer-* like mutant plants (e1-e3) were subjected to sequencing analysis to reveal the indel mutations. **(F)** A chart showing the mutations in the eight *CrRLK1L* genes targeted by CRISPR mutagenesis in line e2 shown in (E). The Sanger sequencing chromatograms are shown in Supplementary Fig. S11.

### Loss of Phosphatidylinositol 4-phosphate 5-kinase 1 (PIP5K1) and PIP5K2 does not significantly affect the function and localization of MLO2

Qin et al. (2020) reported that the loss of PIP5K1 and PIP5K2 significantly impaired MLO2-GFP’s recruitment to the fungal penetration site, resulting in greatly enhanced resistance to an adapted PM fungus (Qin et al., 2020). Based on these findings, they proposed that phosphatidylinositol 4,5-bisphosphate [PI(4,5)P_2_], the product of PIP5K1 and PIP5K2, may be required for MLO2’s focal accumulation, thereby acting as a host susceptibility factor of powdery mildew (Qin et al., 2020). To determine if MLO2’s focal accumulation and function requires PIP5K1 and PIP5K2, we knocked out *PIP5K1* and *PIP5K2* by CRISPR in the *eps3m* line transgenic for *pMLO2*::*MLO2-GFP* and *pRPW8.2::RPW8.2-RFP* and found that the *pip5k1-pip5k2* double mutant plants with greatly reduced stature were also very susceptible to *Gc* UCSC1 (Fig. 10A). Close examination revealed that the first 2-3 true leaves of the mutants were dark purple and often showed reduced infection (Fig. 10B). Sequencing of 10 of 17 independent mutant T1 lines with greatly reduced stature (Supplementary Fig. S15) confirmed the presence of homozygous or biallelic mutations in *PIP5K1* and *PIP5K2*, with the chromatograms of the sequenced mutation sites from two lines (C6 and L16) shown in Fig. 10C. Confocal imaging of infected leaves of the two mutant lines confirmed typical localization of MLO2-GFP at the PPM and normal of RPW8.2-RFP at the EHM (Figure 10D). Collectively, these results demonstrate that PIP5K1 and PIP5K2, and the PI(4,5)P_2_ pool derived from these two enzymes, are dispensable for MLO2’s localization and function.

**Figure 10.**
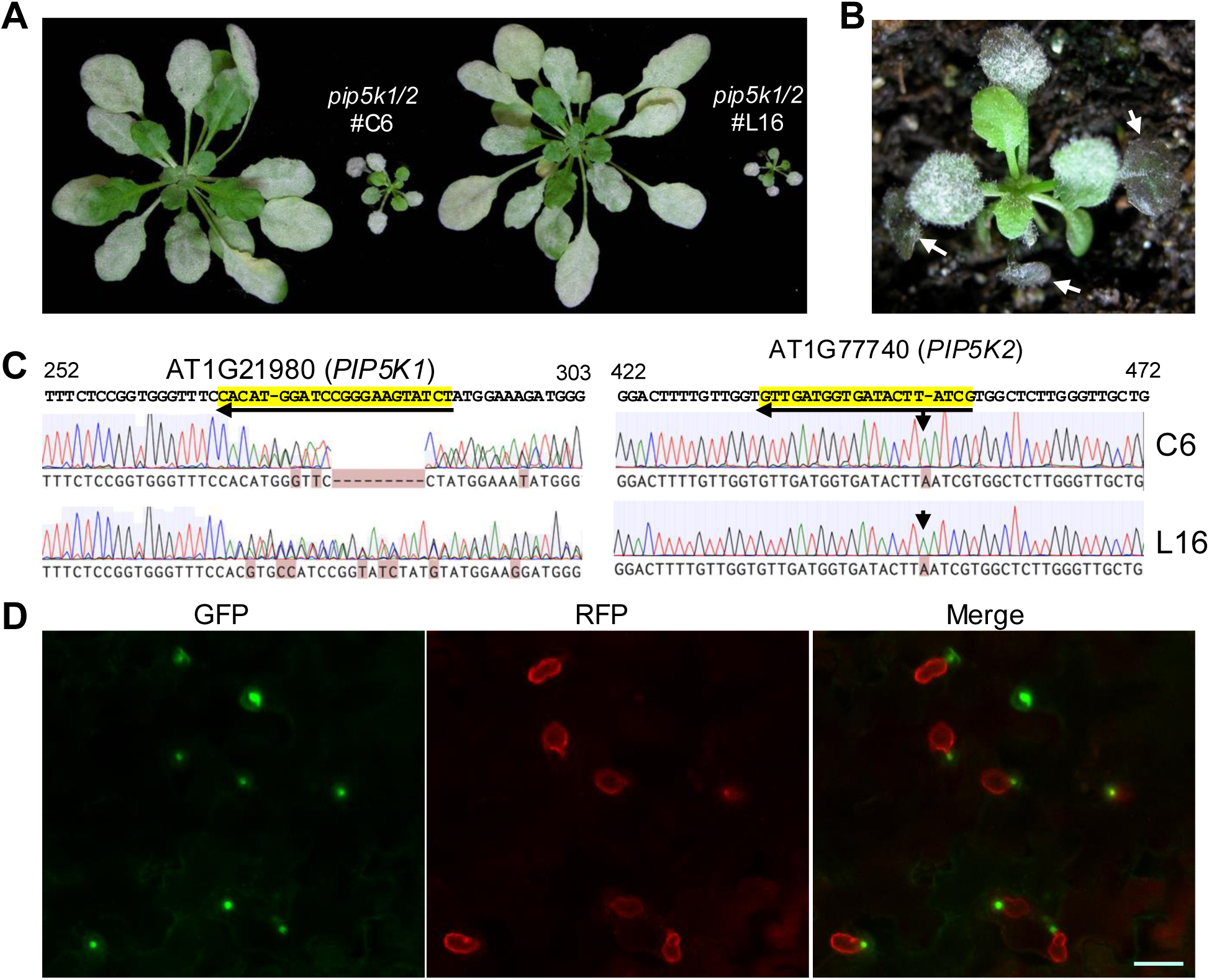
Loss of *PIP5K1/K2* does not affect MLO2’s focal localization and MLO2-mediated susceptibility. CRISPR-targeted mutagenesis was used to knock out *PIP5K1* and *PIP5K2 in eps3m* expressing MLO2-GFP and RPW8.2-RFP. Independent T1 plants were inoculated with *Gc* UCSC1 and plant photos were taken at 10 dpi. Mutations in these two genes in 10 transgenic plants with small stature were identified by targeted sequencing, and two of which were shown in (C). **(A)** Two representative transgenic plants with greatly reduced stature were similarly susceptible as the bigger plants of the parental line. **(B)** A closeup view of a putative *pip5k1/2* mutant plant showing reduced infection of the first 2-3 true leaves (arrows) with anthocyanin accumulation. **(C)** Four chromatographs showing disruptive indels in *PIP5K1* and *PIP5K2* in two selected mutant plants. Long arrowed lines indicate small guide RNAs; short arrows point to the inserted nucleotides. **(D)** A representative confocal image showing normal localization of MLO2-GFP and of RPW8.2-RFP. Bar=20μm.

## Discussion

MLOs have long been considered to be host susceptibility or compatibility factors of powdery mildew (Panstruga, 2003). Yet, to date, there is no definitive genetic evidence to exclude ectopic activation of defense as the molecular basis of *mlo*-mediated “resistance” despite that no specific defense pathways or components have vigorously been shown to contribute to the “resistance” of the *mlo2/mlo6/mlo12* triple mutants (Kuhn et al., 2017). In this study, through a tailored forward genetic screen that aims to identify mutants that display compromised-immunity-yet-poor infection (*cipi*) phenotypes, we discovered disruptive mutations in *MLO2* to be responsible for the *cipi* phenotype of five best *cipi* mutants. CRISPR-targeted mutagenesis of *MLO6* and *MLO12* in *cipi3* and subsequent analyses further demonstrate that *mlo2/6/12*-conditioned suppression of infection can be uncoupled from ectopic activation of plant defenses, providing definitive genetic and molecular evidence to a long-awaited conclusion that MLO proteins are *bona fide* host susceptibility factors that are essential for PM pathogenesis.

### Uncoupling of defense activation from *mlo*-conditioned failure of PM pathogenesis

The concept of host susceptibility factors essential for pathogenesis has long been established in the case of viruses (Strauss and Strauss, 1999). Both RNA and DNA viruses depend on host machinery for their replication, hence mutagenesis of relevant host translation initiation / elongation factors has long been considered effective strategies to engineer viral resistance, particularly in plants (Sanfacon, 2015). A good example of this strategy is provided by a recent report on engineering resistance to maize lethal necrosis caused by maize chlorotic mottle virus along with a potyvirus (Wen et al., 2024). Many “susceptibility” genes have been identified in various plant species that their loss hinders pathogenesis of bacterial, fungal, oomycete pathogens, creating opportunities for engineering plant resistance through targeted mutagenesis (Garcia-Ruiz et al., 2021; Koseoglou et al., 2022). For example, disruption of *DMR6*, which encodes a 2-oxoglutarate (2OG)-Fe(II) oxygenase of previously unknown function, renders Arabidopsis resistant to *Hyaloperonospora parasitica*, an obligate biotrophic oomycete (van Damme et al., 2008). Subsequent studies revealed that loss of *DMR6* homologs leads to salicylic acid (SA)-dependent immune activation and broad-spectrum disease resistance across multiple plant species (Kieu et al., 2021; Thomazella et al., 2021; Tripathi et al., 2021; Giacomelli et al., 2023; Zhang et al., 2025). Thus, *dmr6*-mediated resistance is mostly due to activation of plant immunity. Another group of well-characterized susceptibility genes encode SWEET sugar transporters, which are exploited by *Xanthomonas oryzae pv. oryzae*—the causal agent of rice bacterial blight—through TAL effectors to increase host sugar availability. Natural recessive mutations or CRISPR-enabled editing of *SWEET* genes or their promoters can effectively block sugar exploitation by the bacterial pathogen, thereby conferring resistance to bacterial blight in rice (Chu et al., 2006; Yang et al., 2006; Oliva et al., 2019; Schepler-Luu et al., 2023). The great potential of controlling devasting diseases through identification and targeted interference of susceptibility genes of the causative pathogens was further demonstrated by an elegant study focusing on citrus greening (Zhao et al., 2025). Zhao and colleagues discovered a key susceptibility gene, encoding an E3 ubiquitin ligase, PUB21, and a transcription factor MY2 regulating JA-dependent defense response as the substrate of PUB21. Based this pathogenicity mechanism, the authors searched and identified a 14–amino acid peptide, APP3-14, that can bind and inhibit PUB21 activity, thereby stabilizing MYC2 and ensuring induction of JA-dependent resistance against the causal bacterial pathogen of citrus greening (Zhao et al., 2025). This highlights the importance of understanding the role of host susceptibility factors in pathogenesis.

However, although recessive *mlo* mutation-mediated “resistance” to PM fungi was characterized in 1997 (Buschges et al., 1997) and has been widely exploited in various crop species (Kuhn et al., 2017; Kusch and Panstruga, 2017; Li et al., 2022), its underlying mechanisms remain largely elusive. It is notable that *mlo*-mediated resistance is restricted to clade IV and V almost in all cases [summarized in table 1 in (Li and Xiao, 2025)] even though MLO7, a clade III MLO expressed in synergid cells, can partially complement the loss of MLO2, MO6 and MLO12 in leaf cells (Fig. 7C). Depending on the pathosystems used for study, clade IV and V *mlo* mutants have been shown to be either more susceptible to hemi-biotrophic or necrotrophic pathogens (see a recent review by (Li and Xiao, 2025)). For example, *mlo2/mlo6/mlo12* triple mutants showed increased susceptibility to hemi-biotrophic bacterium *Pseudomonas syringae* (Acevedo-Garcia et al., 2017), necrotrophic fungi *Alternaria alternata* and *A. brasscisicola*, and hemi-biotrophic oomycete *Phytophthora infestans* (Consonni et al., 2006) and *Fusarium oxysporum* (Acevedo-Garcia et al., 2017). However, the Arabidopsis triple mutant exhibited reduced penetration success of hemi-biotrophic *Colletotrichum higginsianum*, which invades host via direct cell wall penetration (Acevedo-Garcia et al., 2017). Similarly, *mlo* mutants of barley, wheat and Medicago showed reduced root colonization by the arbuscular mycorrhizal fungus (Jacott et al., 2020), conforming with the notion that clade IV/V MLOs are required for efficient host cell wall penetration by (hemi)biotrophic filamentous fungi. An exception is that the *m2/6/12* triple mutants showed no altered susceptibility to obligate biotrophic oomycete *Hyaloperonospora arabidopsidis* and *Albugo laibachii* (Acevedo-Garcia et al., 2017)(Fig. 3C). This may be explained by the fact that those oomycetes can enter host mesophyll layers via stomata (Herlihy et al., 2019) and other MLOs expressing in mesophyll cells may serve a similar role in accommodating penetration and haustorium formation of the oomycetes in mesophyll cells. This latter theory can also explain why trichome cells of *eps3m* can still support limited growth of *Gc* UCSC1 (Fig. 4D). Based on the studies discussed above, along with numerous other reports [Reviewed by (Kusch and Panstruga, 2017; Li and Xiao, 2025)], a general pattern emerges regarding the biotic phenotypes resulting from disruption of clade IV/V MLOs: they are consistently associated with compromised biotrophy involving host epidermal cells. In other words, clade IV/V MLOs appear to be essential for the successful penetration of host epidermal cells by biotrophic filamentous fungi—regardless of the nature of the interaction, whether beneficial, commensal, or pathogenic.

The reasoning above is strongly supported by key observations from this study, which was specifically designed to identify genes required for “biotrophy” of PM fungi in host epidermal cells. The isolation of five *mlo2* allelic mutations with the strong *cipi* mutant phenotypes through our genetic screens suggests that MLO2 may be the single most important host protein required for PM pathogenesis (Fig. 1). Consistently, the failure of PM pathogenesis in the *eps3m* mutant could be completely uncoupled from activation of known defenses examined (Fig. 4; Supplementary Fig. S5). Our results also indicate that the early leaf senescence of *3m/C* is due to activation of the EDS1/PAD4 and SA-dependent cellular defense programs, agreeing with earlier inferences (Buschges et al., 1997; Humphry et al., 2006), explaining the enhanced susceptibility of *mlo* mutants to necrotrophic or hemi-biotrophic pathogens (Consonni et al., 2006; Acevedo-Garcia et al., 2017).

### How may MLO2 work to permit PM pathogenesis?

Our genetic and molecular data affirm that MLOs are essential for PM pathogenesis. A critical follow-up question one may ask is what role MLOs play in fulfilling this requirement? Given their obligate biotrophic nature, PM fungi likely co-opt an MLO-dependent host cellular process to achieve successful host penetration and haustorium differentiation. In this context, the accumulation of MLO2-GFP to the PPM, the extracellular membrane domain that matches mostly or exactly the same physical space as the papilla where callose is deposited, or the paramural body where the syntaxin PEN1 (SYP121) exosomes accumulate (Assaad et al., 2004; Bhat et al., 2005) (Meyer et al., 2009), may provide important mechanistic insights. Upon spore inoculation, the earliest focal accumulation of MLO2-GFP was detected at 7.5 hpi (Fig. 6C, D), which is 5-7 hours earlier than the formation of the haustorium underneath the penetration peg, which is estimated to be around 12 to 14 hpi (Koh et al., 2005). Given that germinated sporelings can develop normal appressoria and penetration pegs in *eps3m* (pointed by an arrow in Fig. 3B) but fail to differentiate haustoria in pavement epidermal cells (Supplementary Fig. S4), we speculate that accumulation of functional V MLOs in the PPM may be a prerequisite for haustorial differentiation. More specifically, MLOs may be exocytosed into the PPM and function as scaffolding crucial for stabilizing and resealing the plasma membrane damaged by penetration, which may create a host cell environment for haustorium differentiation from the tip of the penetrate peg (Fig. 6F-J). In addition, a poorly characterized membranous structure, the haustorial neckband (which is indicated as a blue band in Fig. 6J), is thought to form concomitantly with haustorial biogenesis and seal the space between the apoplast and the extra-haustorial membrane matrix (Gil and Gay, 1977; Bushnell and Gay, 1978). Therefore, MLO2 may be also localized to this membrane and required for its formation. Higher-resolution microscopy with immunogold labeling could help confirm this. Furthermore, given MLO2’s calcium channel activity, MLO2-mediated localized calcium influx may be necessary for the resealing and stabilization of the PPM junction including neckband formation, thereby permitting haustorium differentiation inside a host cell.

### Functional diversification of MLOs

MLO1, along with MLO5 and MLO9, is required for pollen tube integrity (Gao et al., 2023), while MLO11 plays a role in root thigmomorphogenesis (Chen et al., 2009). Results from our analyses through transgene expression using the *MLO2* promoter demonstrate that there is clear functional diversification among MLO1 (clade II), MLO11 (clade I), MLO2, MLO6 (clade V) and MLO7 (clade III) both in terms of subcellular localization and their ability in permitting PM pathogenesis (Fig. 7). While MLO2, MLO6 and MLO7 were also found in the plasma membrane of epidermal cells (Fig. 6E and Fig. 7B,D), they mainly exhibit punctum distribution and relocalize to the PPM in PM-infected cells to accommodate PM pathogenesis. In contrast, MLO1 and MLO11 are homogenously localized at the plasma membrane, show no obvious focal accumulation at the PPM, and cannot substitute MLO2 in permitting PM pathogenesis (Fig. 7F,H). Given that MLO1, MLO2, and MLO7 all exhibit calcium channel activity (Gao et al., 2022; Gao et al., 2023), their functional diversification may stem from their regulation by different CaMs or CMLs or other proteins, rather than from their ability to conduct calcium ion across the plasma membrane. Alternatively, they do not possess calcium channel activity in epidermal cells and/or such activity is not important for their biological functions.

Results from domain swapping between MLO1 and MLO2 (Fig. 8) and between MLO1 and MLO7 (Jones et al., 2017) suggest that there are at least two functional arms for MLO proteins: the N-terminal 7TM portion performs the cellular function of a particular MLO protein whereas the C-terminus specifies its subcellular localization. Such a functional configuration can explain the functionality of chimeric proteins MLO2n-1c in conferring partial susceptibility of *eps3m* to *Gc* UCSC1 (Fig. 8F,G) and NTA (MLO7)-MLO1^Cterm^ (also named faNTA) in restoration of fertility of *mlo7* (Ju et al., 2021). Notably, because MLO2n-1c does not show significant focal accumulation, its presence in the PPM is likely limited. This probably explains the weak susceptibility of *eps3m* lines expressing MLO2n-1c to *Gc* UCSC1, underscoring the importance of a “dosage” effect through MLO2’s C-terminal domain-mediated focal accumulation. This also aligns with the observed correlation between MLO2 expression levels and the susceptibility of transgenic lines (Fig. 5). On the other hand, MLO1n-2c exhibited punctum distribution and focal accumulation induced by sporelings in *eps* (Fig. 8E), but it did not confer susceptibility in *eps3m* (Fig. 8B,C), supporting the role of the C-terminus of MLO2 in focal accumulation.

It is also worth noting that MLO10, MLO7’s closest family member, displayed punctum distribution (Fig. 7J,L) but only allowed very limited PM growth in the best-case scenario (Fig. 7K). In synergid cells, MLO10, but not MLO8, could restore fertility of *mlo7* (Jones et al., 2017), indicating functional diversification among these three clade III family members. Given this complexity, and the lack of localization data for MLO3, MLO4 and MLO5 in Arabidopsis leaf epidermal cells, it is difficult to infer which sequence features or polymorphisms underlie MLOs’ contrasting molecular functions as reflected by permitting PM pathogenesis and displaying signal-induced relocalization.

### Regulation of MLO2’s focal accumulation at the PPM

Regarding PM penetration-induced MLO2 relocalization to the PPM (Fig. 6), a key question is what specific signal(s) triggers its polarizing trafficking and how this process is regulated. Given that MLO2’s CaMBD-containing cytoplasmic C-terminus directs its focal accumulation, it is conceivable that changes in cytosolic calcium concentration—triggered by calcium influxes during fungal penetration of the host cell wall or plasma membrane—could induce MLO2-calmodulin binding or dissociation, thereby triggering MLO2’s relocalization to the PPM. Such a mechanism would imply that changes of cytosolic calcium levels differentially impact MLOs residing in different subcellular compartments (e.g., MLO2 mostly at the ER/Golgi versus MLO1 at the plasma membrane), likely due to structural differences in their C-termini and specific calmodulins they interact with. In this context, it is worth noting that MLO1n-2c remained its punctum distribution in leaf epidermal cells of *eps3m* insulted by sporelings (Fig. 8C). However, MLO1n-2c were relocalized to the PPM in *eps* (Fig. 8G), suggesting that host cell-wall or plasma membrane disruption induced by successful host entry triggers PPM-oriented trafficking pathway, resulting in dispatch of ER/Golgi-localized MLO2 and the non-functional MLO1n-2c to the PPM. Because successful host entry requires functional MLO2, this would imply a positive feedback mechanism for MLO2’s focal accumulation at the PPM. Without functional plasma membrane-localized MLO2 to initiate the ER/Golgi→ PPM-directed trafficking, MLO1n-2c would remain sequestered at the ER/Golgi.

How MLO2’s ER/Golgi → PPM trafficking polarity is established remains unknown. Phosphatidylinositol-4,5-biphosphate [PI(4,5)P_2_] is a lipid messenger that plays crucial role in polarized trafficking, guiding vesicle and protein movement to specific subcellular locations in both animals and plants (Mei et al., 2012; Thapa and Anderson, 2012). Qin et al. reported that disruptive mutations in *PIP5K1* and *PIP5K2* significantly reduced MLO2-GFP’s focal accumulation at the fungal penetrations site (Qin et al., 2020). Because PIP5Ks catalyze the phosphorylation of phosphatidylinositol 4-phosphate [PI(4)P] to produce PI(4,5)P_2_, this observation suggests that PI(4,5)P_2_ may serve as a trafficking cue in directing MLO2 to the PPM. Unfortunately, results from our genetic study showed that both MLO2-GFP’s focal accumulation at the PPM and MLO2-mediated susceptibility to *Gc* UCSC1 remained unchanged in the absence of PIP5K1 and PIP5K2 (Fig. 10). This suggests that PI(4,5)P_2_ unlikely serves as a trafficking cue for MLO2’s enrichment at the PPM.

FER is required for MLO7’s relocalization from Golgi bodies to the filiform apparatus in synergid cells (Jones et al., 2017; Ju et al., 2021). However, our genetic data indicate that FER and probably five other CrRLK1L family members are dispensable for MLO2’s focal accumulation and its-mediated PM pathogenesis (Fig. 9). This suggests that either there is deeper functional redundancy among the 17 CrRLK1L family members (Lindner et al., 2012) or a CrRLK1L-independent mechanism distinct from that employed in synergid cells for relocalization of MLO7 is engaged for the regulation of MLO2’s re-localization to the PPM in leaf epidermal cells.

In conclusion, data from this study definitively demonstrate that MLO2, MLO6, MLO12 are *bona fide* susceptibility factors of PM fungi. MLO2 (and, by inference, clade IV and V MLOs) appears to adopt a bipartite functional configuration: while the N-terminal 7TM portion executes a specific cellular function, such as cell wall and/or plasma membrane sealing and stabilization, the CaMBD-containing C-terminus orchestrates its spatiotemporal activity via signal-induced re-localization and enrichment. Additionally, our demonstration of the protein dosage effect for MLO2 in mediating susceptibility points to the potential for engineering PM “resistance” without severe early leaf senescence by downregulation of target MLOs via virus-induced gene silencing (VIGS) or promoter editing. Future investigations will focus on elucidating how clade V MLOs contribute to the sealing and stabilization of the plasma membrane–penetration peg–haustorial neckband junction, thereby accommodating haustorium differentiation and PM pathogenesis.

## Materials and Methods

### Plant lines and growth conditions

All mutants used in this study were in the *Arabidopsis thaliana* accession Col-0 background. Mutants *eds1-2* (Bartsch et al., 2006), *pad4-1* (Jirage et al., 1999) and *sid2-2* (Wildermuth et al., 2001) have been previously described. The triple mutant *eds1-2/pad4-1/sid2-2* mutant was generated by genetic crosses and identified by PCR genotyping as previously described (Zhang et al., 2018). The *mlo2-5/mlo6-2/mlo12-1* triple mutant was previously described (Consonni et al., 2006). Seeds were sown in SunGro Horticulture (Agawam, MA, U.S.) and cold treated (4 ℃ for 2 days) before moving to growth chambers. Seedlings were transplanted and kept growing under 22 ℃, 75% relative humidity, short day (8 h light at ∼125 μmol m-2 s-1, 16 h dark) conditions for up to 14 weeks before use.

### EMS mutagenesis of seeds and mutant screening

About 10,000 eps seeds (approx. 200 mg) were placed in a 250 ml glass flask. 15 mL dH2O and 30 μL 0.2% EMS were added and the seeds were shaken overnight. After washing with dH_2_O, seeds were blotted dry and mixed with 250 g fine sand and aliquoted into about 50 parts. Each aliquot of ∼5 g sand with seeds was sown evenly in SunGro Horticulture (Agawam, MA, U.S.) in 50 flats. Seedling were grown in a greenhouse for about two months till maturity. Seeds from 30-90 M1 plants were collected to make one M1 seed pool. A total of 102 M1 seed pools were obtained. For mutant screening, about 150-450 M2 plants per pool were prepared in one or two flats for inoculation with *Gc* UCSC1. Putative *cipi* mutants were transplanted into individual pots for further growth till maturity.

### Pathogen Infection, Disease Phenotyping, and Quantification

The Arabidopsis-adapted powdery mildew isolate *Golovinomyces cichoracearum* (*Gc*) UCSC1 was maintained on live Col-0 or *eds1-2* plants. Inoculation, visual scoring of disease reaction phenotypes and spore quantification were done as previously described (Xiao et al., 2005).

### Mapping of causal mutations

Bulked segregant pool-genome sequencing was used to identify candidate causal mutations. *cipi2* and *cipi3* mutants were crossed with the *eps* parental line, and their corresponding F2 segregating populations were inoculated with *Gc* UCSC1. Genomic DNA was isolated from the mixture of the leaf tissues harvested from 65 or more F2 individuals showing the *cipi* phenotypes using NucleoSpin Plant II kit (MACHEREY-NAGEL, #740770.50). About 2 μg of each DNA sample was sent to BGI (Beijing Genomics Institute, Shenzhen, China) for deep sequencing using Illumina Hiseq 2000 platform at an average coverage of ∼50x with 100 bp paired end reads. The subsequent sequence analysis was done according to the method previously described (Wang et al., 2018). Briefly, sequencing reads were mapped against the TAIR10 Arabidopsis reference genome using Bowtie (Langmead et al., 2009) and variants were called by SAMtools (Li et al., 2009). Only G to A and C to T conversions predominantly caused by EMS mutagenesis were picked for further analysis. The effect of each EMS-induced SNP (single-nucleotide polymorphism) on the corresponding gene was annotated using snpEffect (Cingolani et al., 2012). The effects of the SNPs were classified into three categories: very high (stop gained, splice site donor, splice site acceptor), high (non-synonymous coding, start gained, stop lost), and other (intergenic, intron, 3’ UTR, 5’ UTR, synonymous coding).

### DNA constructs and Arabidopsis transformation

The pK7FWG2 plasmid (Karimi et al., 2002) was used for cloning of all *MLO* genes in translational fusion with eGFP. To replace the original *35S* promoter with the *MLO2* native promoter (pMLO2), restriction enzymes *Xba*I and *Bam*HI were used to linearize the plasmid, and ∼2 Kb *pMLO2* fragment amplified from genomic DNA was inserted, leading to a *pMLO2::ccdB-eGFP* cassette used for cloning *MLOx-eGFP* fusion genes under control of the *MLO2* promoter. To create various *MLO* expression constructs, corresponding *MLO* genes were first cloned into the pENTR/D-TOPO vector via TOPO cloning (https://www.thermofisher.com/us/en/home/life-science/cloning/topo.html). Then, the *MLO* genes were shuttled to the binary vector containing *pMLO2::ccdB-eGFP* via Gateway LR reaction. All vectors are verified by Sanger sequencing.

Amplification of fragments for creation of chimeric *MLOs* was done using primers in Supplementary Table S1 by overlap-extension PCR. Specifically, the two fragments were amplified from Col-0 cDNA with overlapped chimeric primers. The two products were then mixed together at the same molar concentration and served as template for amplification of the full-length chimeric gene. All vectors are verified by Sanger sequencing.

All binary vectors containing the *MLOx-eGFP* constructs were transfected into *A. tumefaciens* GV3101. Arabidopsis transformation was conducted following the floral dipping protocol described previously (Clough and Bent, 1998).

### Agrobacterium-mediated transient expression in *Nicotiana benthamiana* leaves

*N. benthamiana* plants were grown in a growth chamber under 22 ℃, 75% relative humidity, long day (16 h light at ∼125 μmol m^−2^ s^−1^, 8h dark) conditions for four weeks, and then were moved to short day condition (8 h light at ∼125 μmol m^−2^ s^−1^, 16h dark) for 1 week before agroinfiltration. *A. tumefaciens* GV3101 cells containing corresponding vector(s) were suspended in infiltration buffer [10 mM MgCl_2_, 10 mM MES (pH 5.6), and 200 μM acetosyringone] to a final OD_600_ value of 0.4-0.6. The agrobacterium suspension was incubated in dark for 2 h before infiltration into *N. benthamiana* leaves using a blunt syringe. The agroinfiltrated leaves were examined for MLOx-eGFP expression by confocal microscopy following the method described below.

### Confocal microscopy

The expression and localization of the MLOx-eGFP fusion proteins were examined by confocal microscopy using a Zeiss LSM710 microscope. Saturated pixels are intentionally avoided. Confocal images were post-processed using ZEN software (Carl Zeiss, 2009 edition) with global linear adjustments applied consistently and in accordance with academic best practices for image processing.

### Detection of H_2_O_2_ Accumulation and cell death

DAB (3,3’-diaminobenzidine) staining was used to detect *in situ* H_2_O_2_ production and accumulation while trypan blue staining was used to detect dead or dying cells as well as fungal structures in leaves. These methods were previously described (Xiao et al., 2003).

### Quantitative RT-qPCR analysis

Three leaf samples of seven-week-old plants (∼100 mg) per genotype were harvested before and at 0 hpi, 6 hpi, 12 hpi and 48 hpi after *Gc* UCSC1 infection. Total RNA was extracted using TRIzol® Reagent and reverse transcribed into cDNA using SuperScriptTM III Reverse Transcriptase (Invitrogen, Thermo Fisher Scientific Inc.). For each experiment, qRT-PCR was performed with three biological replicates per treatment and three technical replicates per sample using the Applied Biosystems 7300 Real-Time PCR System with SYBRTM Green PCR Master Mix (Thermo Fisher Scientific Inc.). The transcript levels of the target genes were normalized to that of *UBC9* (Ubiquitin conjugating enzyme 9, AT4G27960). Data was analyzed using the Applied Biosystems 7300 Real-Time PCR System Software and the comparative ΔCt method. Primers used for qRT-PCR are listed in Supplementary Table S2.

### Western blot

Leaf tissue of 150 mg per sample was frozen in liquid nitrogen and grounded into fine powder. Two volume (w/v) Ripa buffer containing 100 μM PMSF, 1 x protease inhibitor, and 100 mM DTT was added to the frozen sample and then vortexed for homogenization. Samples were incubated at 95 ℃ for 5 min in 1 x SDS sample buffer, then centrifuged at 12000 g for 10 min. Samples were loaded to 4-12% Bis-Tris SurePAGE™ gel (GenScript #M00652) for electrophoresis following manufacturer’s instruction. Membrane transfer was conducted by 100 V for 2 h. The membrane was blocked with 5% BSA in TBST for 1h and then incubated with anti-GFP antibody (ab290, Abcam) overnight at 4 ℃. After 3×10 min washing with TBST, the membrane was loaded with secondary antibody (and incubated at room temperature for 1 h with shaking, followed by 3×10 min wash. The signal was generated by Clarity Western ECL Substrate (BIO-RAD ##1705061) and imaged with BIO-RAD ChemiDoc Imaging System.

### CRISPR/Cas9 targeted mutagenesis

Two CRISPR/Cas9 genome editing systems were used for targeted mutagenesis of genes of interest in this study and all related plasmids were purchased from Addgene (https://www.addgene.org). The first utilizes an Arabidopsis egg cell-specific promoter to drive the expression of Cas9 (Wang et al., 2015). This system was used to knockout *MLO2*, *MLO6* and *MLO12*. The other system was developed for multiplexed CRISPR (Stuttmann et al., 2021). It utilizes an intronized Cas9 to improve Cas9 expression and editing efficiency. This system was used to knockout *FER* and its other *CrRLKL1* family members, as well as *PIP5K1* and *PIP5K2*. The pDGE347 binary destination vector was used for cloning the guide RNA constructs. All recombinant plasmids containing the guide RNA cassettes were confirmed by sequencing. All guide RNA sequences were listed in Supplementary Table S3.

### Genotyping of mutants and transgenic lines

All primers used for genotyping are listed in Supplementary Table S4. To detect the *mlo2* allele from *cipi2*, the fragment containing the mutation was amplified by MLO2-e9F/MLO2-e11R followed by *Mwo*I digestion (New England Biolabs, R0573S). The wild type allele is cut into two smaller fragments (161+242 bp), while the *cipi2* mutant allele is intact (403 bp). Similarly, to detect the *mlo2* allele from *cipi3*, the fragment containing the mutation was amplified by MLO2-e2F/MLO2-e3R followed by *Hind*III digestion (New England Biolabs, R3104S). While the wild type allele is intact (341 bp), the *cipi3* mutant allele is digested into two fragments (179+162 bp). The remaining *mlo2* alleles *cipi11*, *cipi12*, and *cipi15* were amplified with MLO2e2F/MLO2-e3R, MLO2-e1F/MLO2-e3R, and MLO2-e6F/MLO2-e8R, respectively, followed by sequencing to detect the respective mutations. To detect mutations in *FER* and its family members, and in *PIP5K1* and *PIP5K2*, the guide RNA target regions were amplified with gene-specific primers (listed in Supplementary Table S4). PCR products were Sanger-sequenced. Benchling (https://benchling.com/), a cloud-based platform for molecular biology data analysis, was used to make sequence alignments for identification of indels.

### Accession numbers

DNA Sequences of the genes studied in this study can be found in the National Center for Biotechnology Information (NCBI) or The Arabidopsis Information Resource under the following gene ID numbers: AT4G02600 (Arabidopsis *MLO1*), AT1G11310 (Arabidopsis *MLO2*), AT3G45290 (Arabidopsis *MLO3*), AT1G11000 (Arabidopsis *MLO4*), AT2G33670 (Arabidopsis *MLO5*), AT1G61560 (Arabidopsis *MLO6*), AT2G17430 (Arabidopsis *MLO7*), AT5G65970 (Arabidopsis *MLO10*), AT5G53760 (Arabidopsis *MLO11*), AT2G39200 (Arabidopsis *MLO12*), AT2G19190 (Arabidopsis *FRK1*), AT5G44420 (Arabidopsis *PDF1.2*), AT5G14930 (Arabidopsis *SAG101*), AT2G14610 (Arabidopsis *PR1*), At1g21980 (Arabidopsis *PIP5K1*), At1g77740 (Arabidopsis *PIP5K2*), At5g38990 (Arabidopsis *MDS1*), At5g59700 (Arabidopsis *ANJ*), At5g54380 (Arabidopsis *THE1*), At1g30570 (Arabidopsis *HERK2*), At3g51550 (Arabidopsis *FER*), and unmade CrRLK1L family members, At2g23200, At2g39360, and At5g24010.

## Supplementary data

**Supplementary Figure S1.**
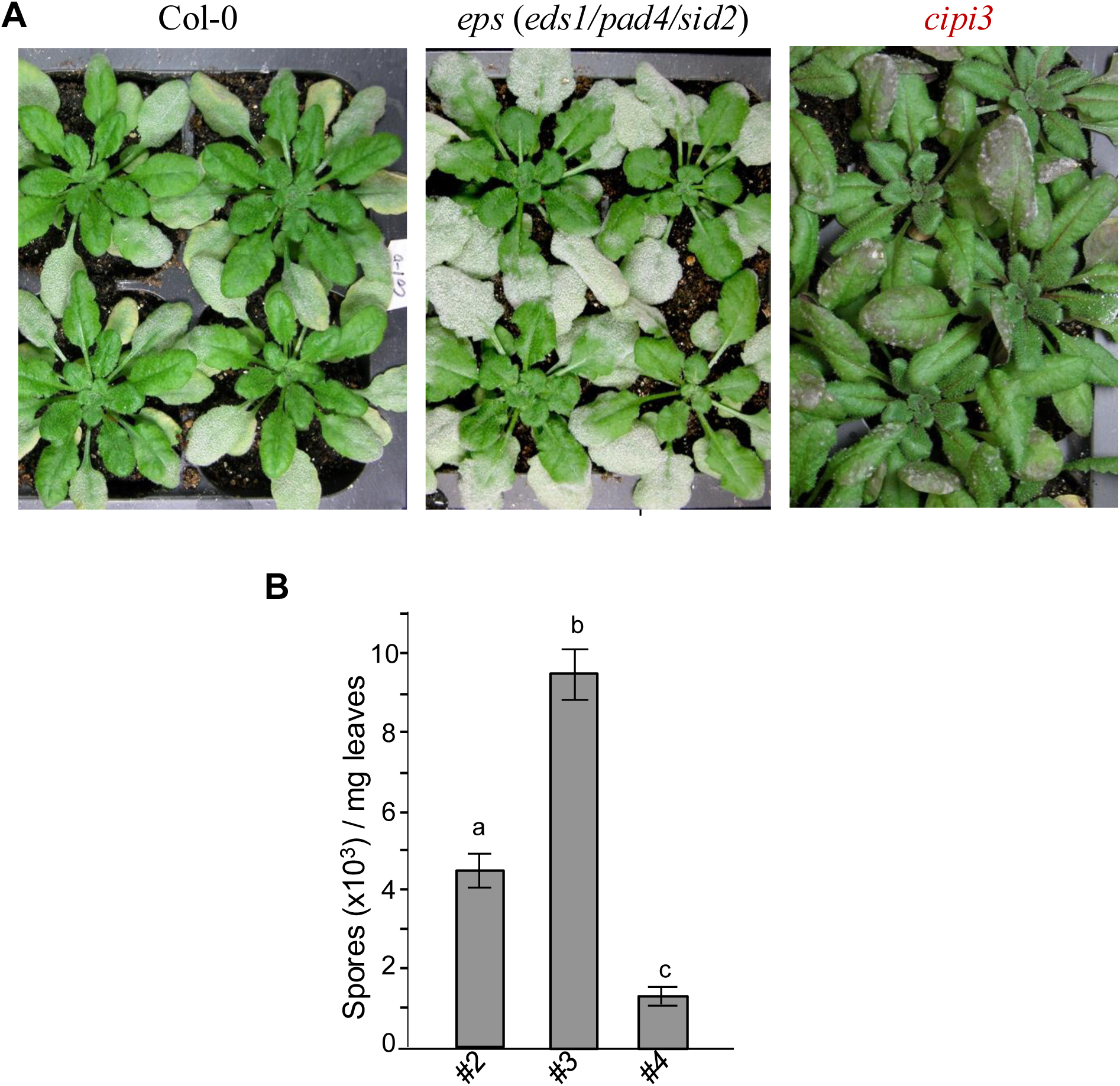
*cipi3* exhibits strong “resistance” to *Golovinomyces cichoracearum* (*Gc*) UCSC1. **(A)** Eight-week-old plants of the indicated genotypes were infected with *Gc* UCSC1. Photos were taken at 10 dpi. Note the trichome-based infection in *cipi3*. **(B)** Reduced susceptibility of *cipi3* compared to Col-0 wild-type and the *eps* (*eds1/pad4/sid2*) parental line as measure by total number of spores per mg infected leaves at 12 dpi. Different letters indicate statistically significant differences (*P*<0.001) between the three lines, as determined by multiple comparisons using one-way ANOVA, followed by Tukey’s HSD test. This experiment was repeated once with similar results.

**Supplementary Figure S2.**
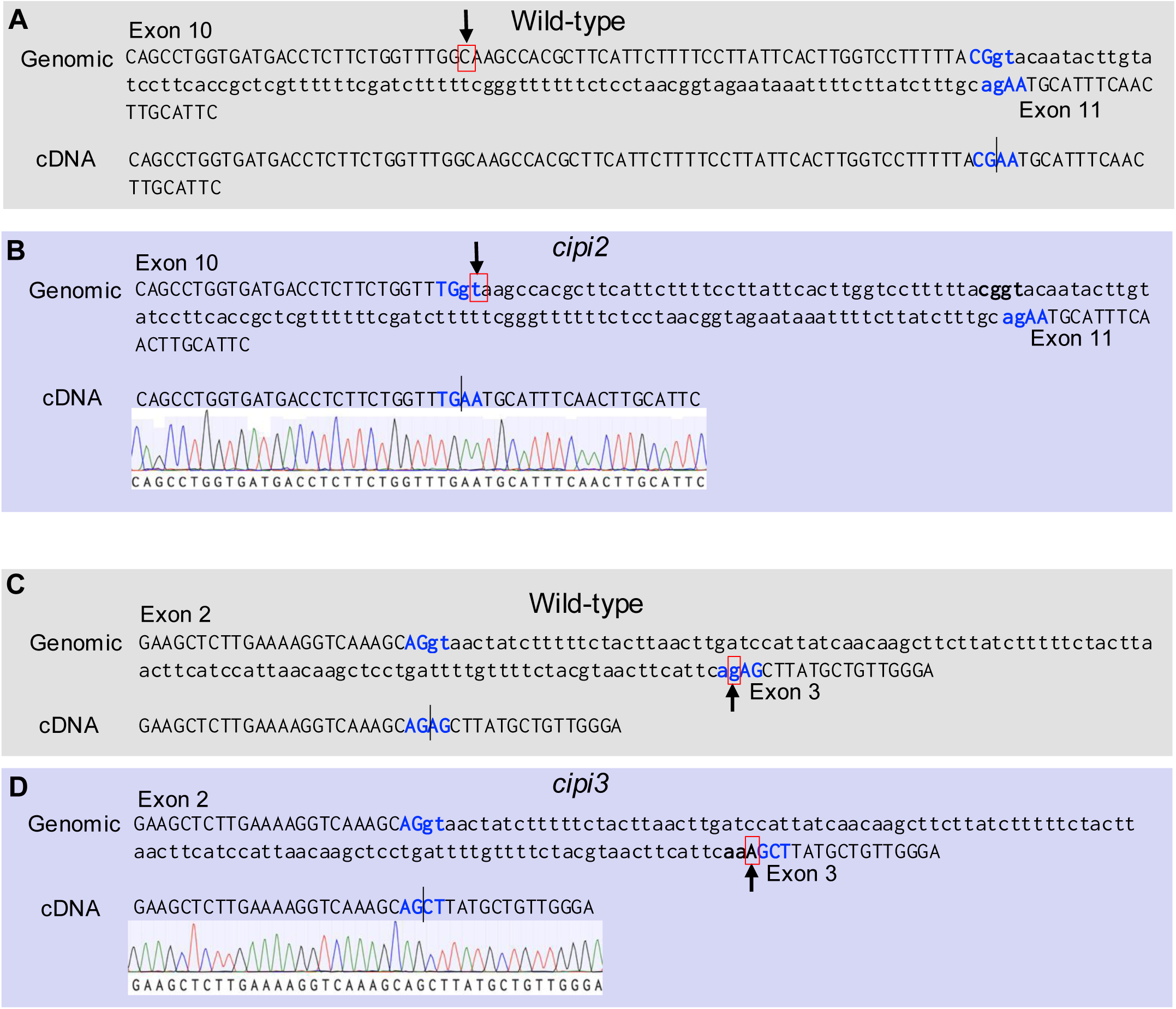
Predicted and sequence-confirmed mis-splicing of *MLO2* mRNA in the *cipi2* and *cipi3* mutants. Shown are the exon-intron boundaries and the genomic DNA (**A,C**) and predicted and sequence-confirmed cDNAs (**B,D**) reflecting mRNAs resulted from mRNA splicing in wild-type and the mutation regions of *cipi2* (**A,B**) and *cipi3* (**C,D**). The nucleotides of the introns are in lower case. The splicing motifs are in highlighted in blue. The vertical bar indicates the splice boundary after removal of the intron in the mRNA. Arrows indicate the C -to-T mutation in *cipi2* (**B**) or G-to-A the mutation in *cipi3* (**D**) confirmed by the respective Sanger sequencing chromatographs.

**Supplementary Figure S3.**
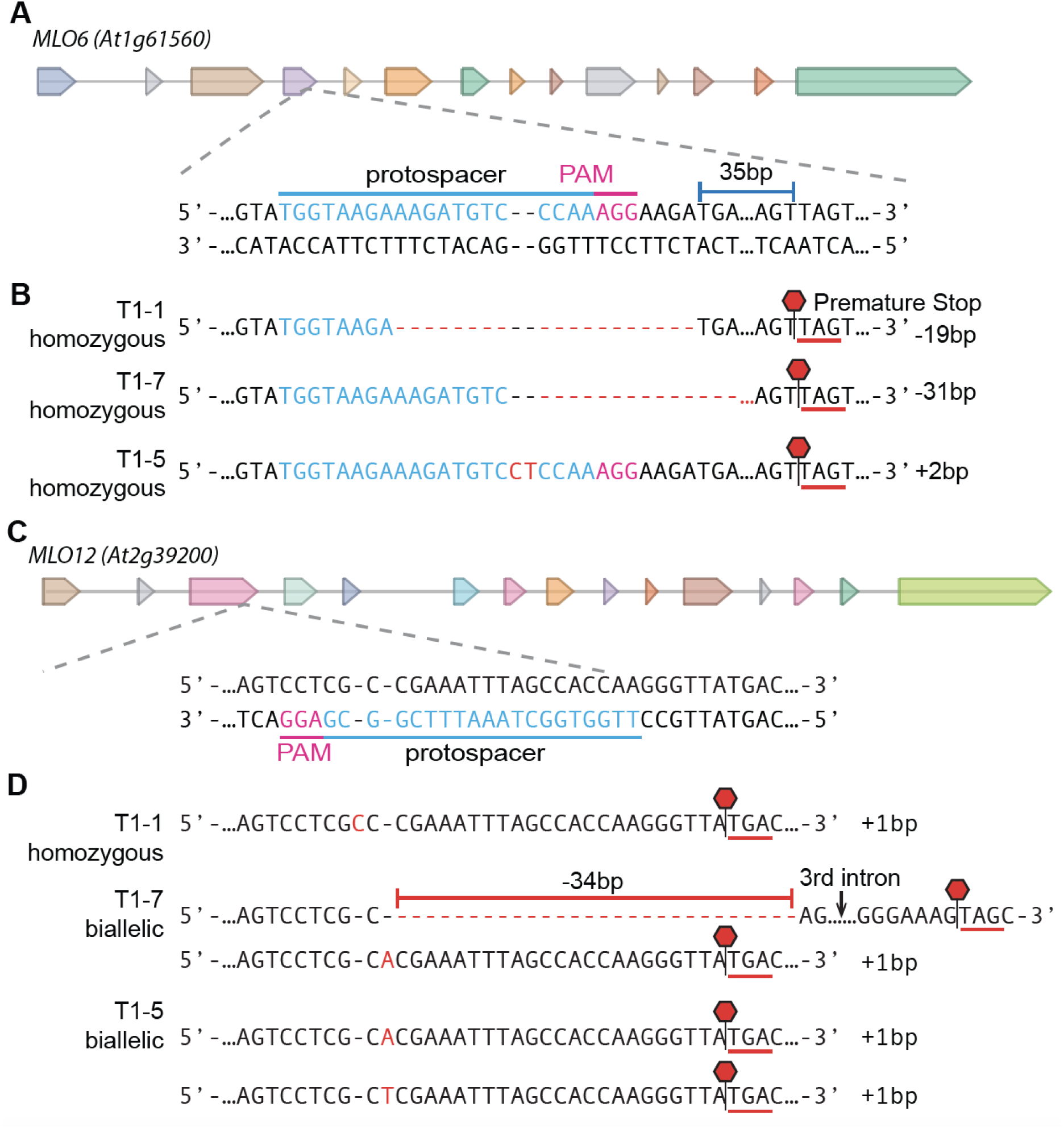
CRISPR-targeted mutagenesis of *MLO6* and *MLO12*. **(A)** MLO6’s gene structure with the protospacer and PAM sequence marked. **(B)** Three independent lines with indels in *MLO6* and the position of the premature stop codon marked with red hexagons. **(C)** MLO12’s gene structure with the protospacer and PAM sequence marked. **(D)** Three independent lines with indels in *MLO12* and the position of premature stop codon marked with red hexagons.

**Supplementary Figure S4.**
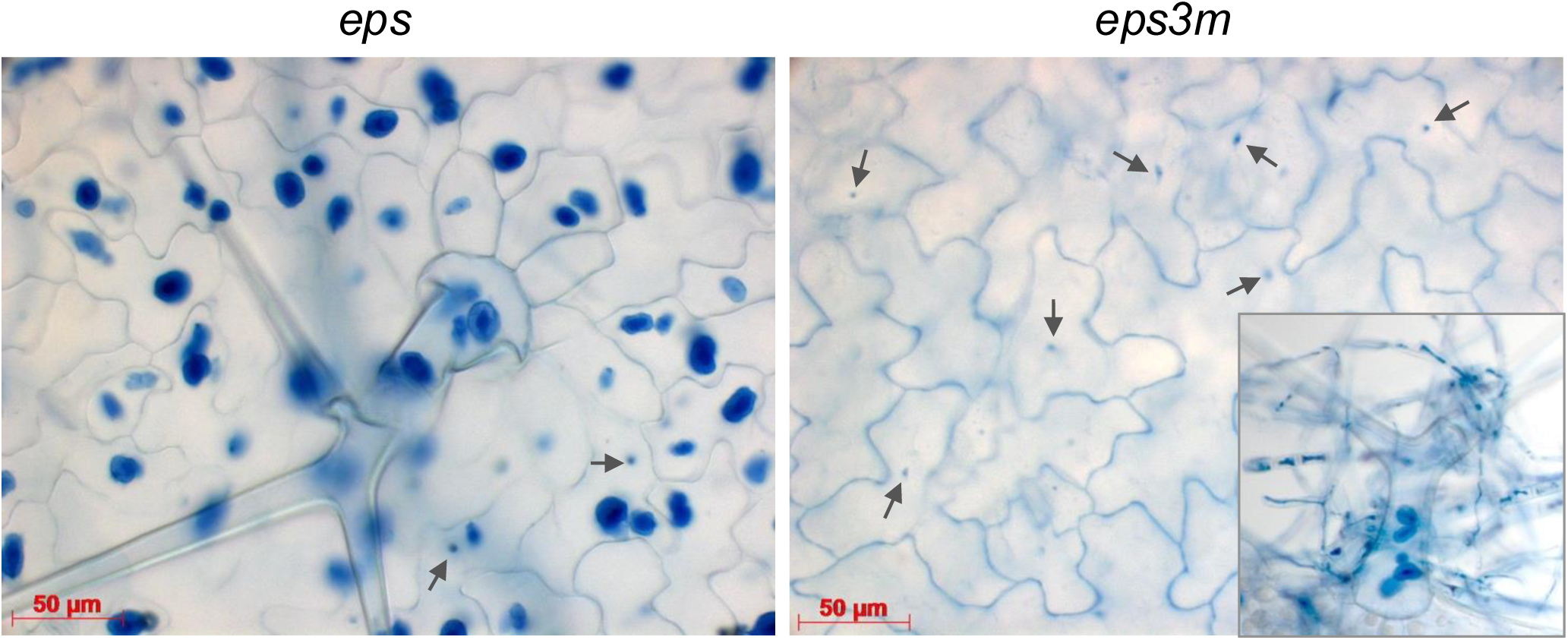
No haustorium formation in pavement cells of *eps3m*. Plants of *eps* and *eps3m* were inoculated with *Gc* UCSC1. At 6 dpi, infected leaves were gently brushed using a fine brush under tap water for 30 seconds to completely remove fungal sporelings and mycelia on the leaf surface and then subjected to trypan blue staining. Dark blue-stained haustoria are visible in the pavement cells of *eps* but not found in those of *eps3m*. Instead, small blue dots (arrows) likely representing penetration sites were visible in pavement cells of *eps3m*. In contrast, haustoria can be found in infected trichome cells of *eps* and *eps3m* (inset).

**Supplementary Figure S5.**
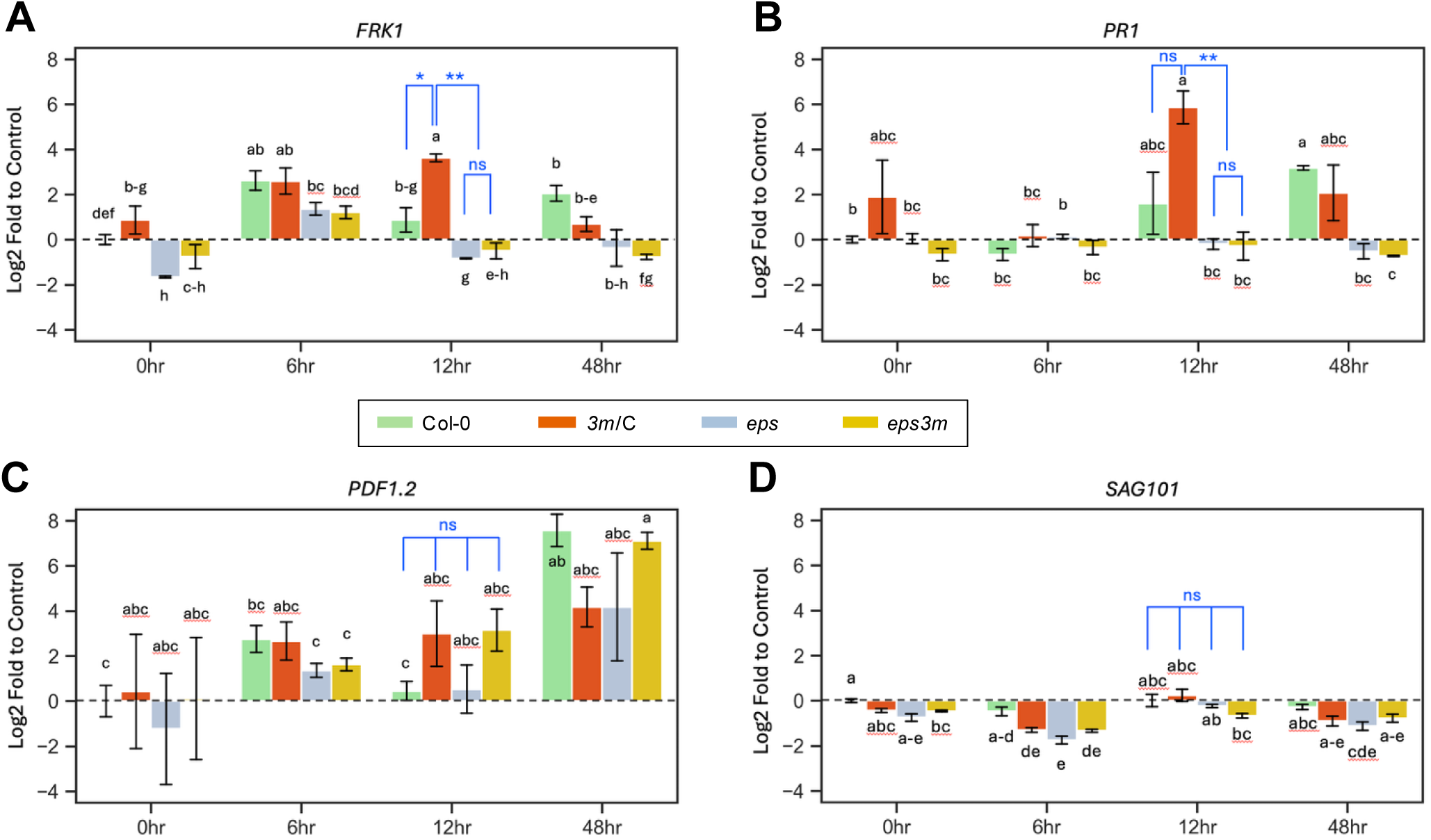
No activation of PTI and ETI marker genes in *eps3m*. Six-week-old, short-day grown plants of the four indicated genotypes were inoculated with *Gc* UCSC1. Inoculated leaves were collected at the indicated four timepoints and subjected to qRT-qPCR analysis to measure expression of four indicated marker genes. The control group is defined as Col-0, 0h. All gene groups pass ANOVA with *p*<0.05. Post-hoc analyses (multiple comparisons) are conducted through *T*-test adjusted by Benjamini-Hochberg FDR procedure. \**p*<0.05, \*\**p*<0.01, \*\*\**p*<0.001”. **(A)** Expression of *FRK1*, reporting activation of PTI. **(B)** Expression of *PR1*, reporting activation of ETI. **(C)** Expression of *PDF1.2*, reporting activation of the Jasmonic acid- and ethylene-dependent defenses. **(D)** Expression of *SAG101*, reporting the onset of leaf senescence. This experiment was repeated twice with similar results.

**Supplementary Figure S6.**
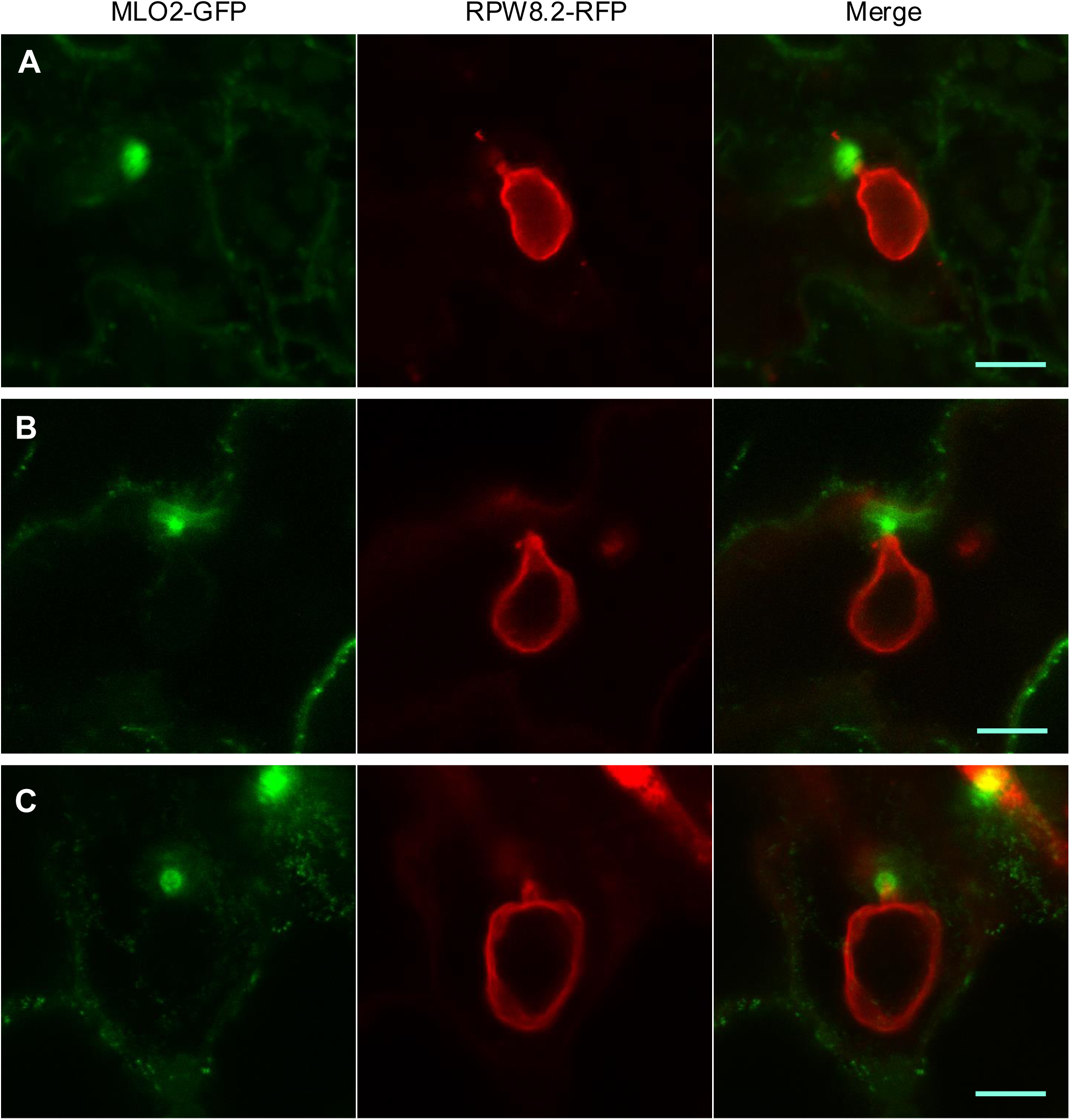
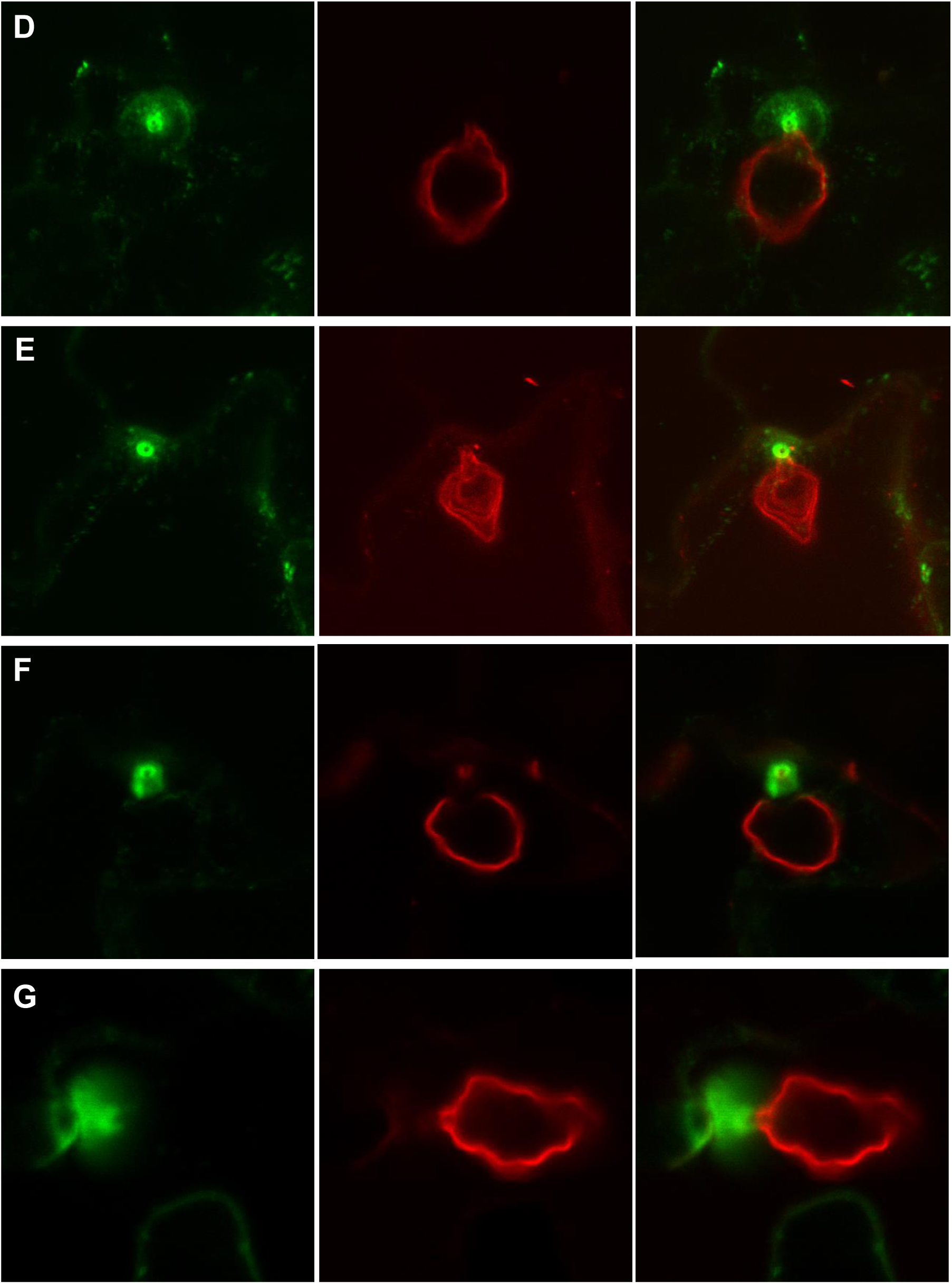
MLO2-GFP is localized to the peri-penetration peg membranous space (PPM) next to the haustorial neck. Plants of *eps3m* expressing MLO2-GFP and RPW8.2-RFP were inoculated with *Gc* UCSC1. Infected leaves were subjected to confocal microscopy at 2 -3 dpi. **(A-C)** Three z-stack (3-5) projected confocal images with the GFP and RFP individual channels. Note, the merged images are shown in Fig. 7F**-H**. Bar=10μm. **(D-G)** Additional four confocal images from 3-5 z-stack projection. Bar=10μm.

**Supplementary Figure S7.**
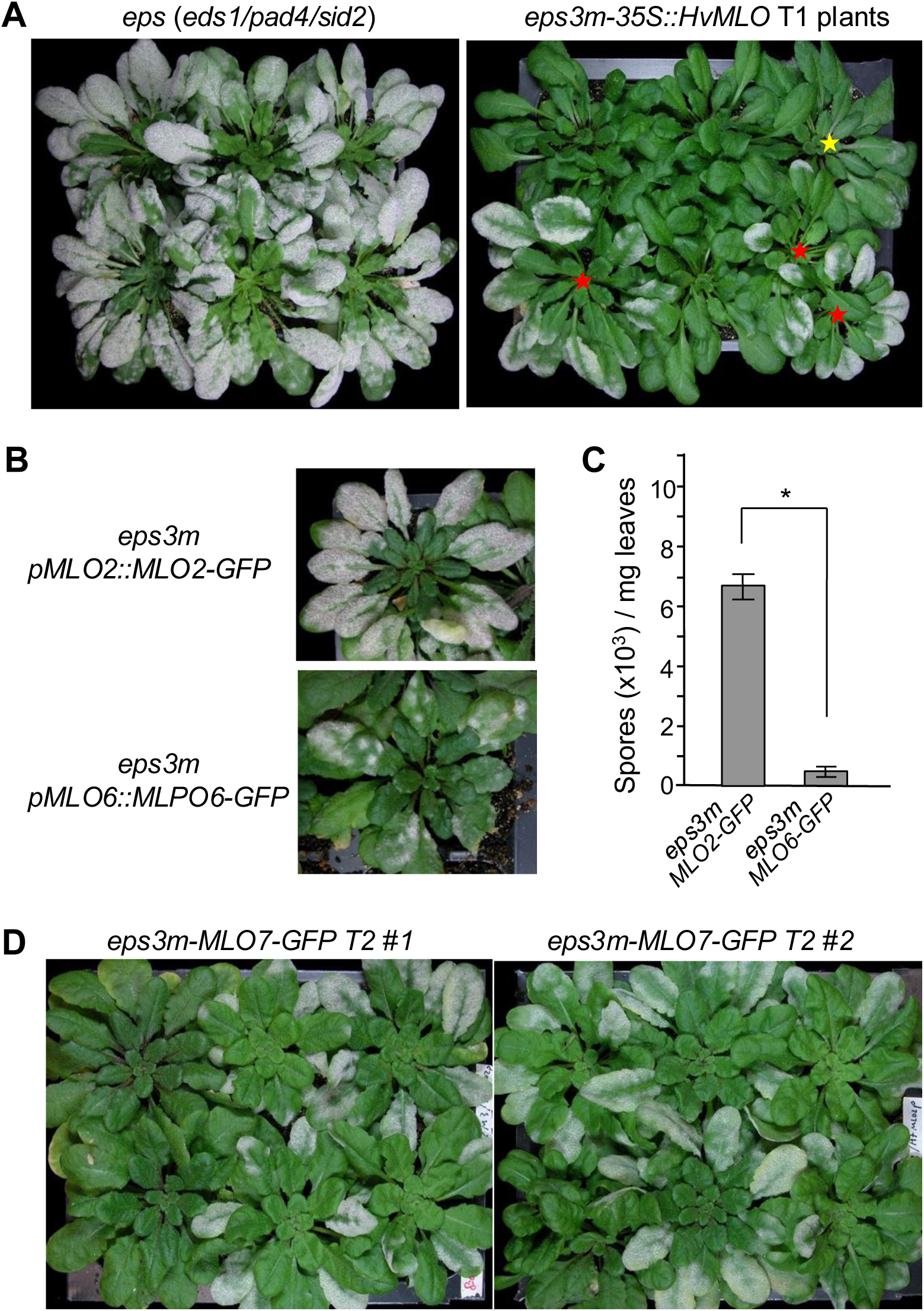
Expression of HvMLO1-GFP, MLO6-GFP or MLO7-GFP in *eps3m* partially restored susceptibility to *Gc* UCSC1. **(A)** Representative T1 plants of *eps3m* transgenic for *35S::HvMLO1* infected with *Gc* UCSC1 at 12 dpi. Note, three T1 plants were moderately susceptible (red stars) while one was weakly susceptible (yellow star) compared to *eps* plants. **(B,C)** Representative photos of the indicated transgenic lines infected with *Gc* UCSC1 at 11 dpi (B) and their levels of susceptibility (C). Asterisk indicates significant difference (*p*<0.001; unpaired Student’s *t*-test). **(D)** Infection phenotypes of the T2 progenies of two *eps3m* lines transgenic for *pMLO2::MLO7-GFP*. Photos were taken at 11 dpi with *Gc* UCSC1.

**Supplementary Figure S8.**
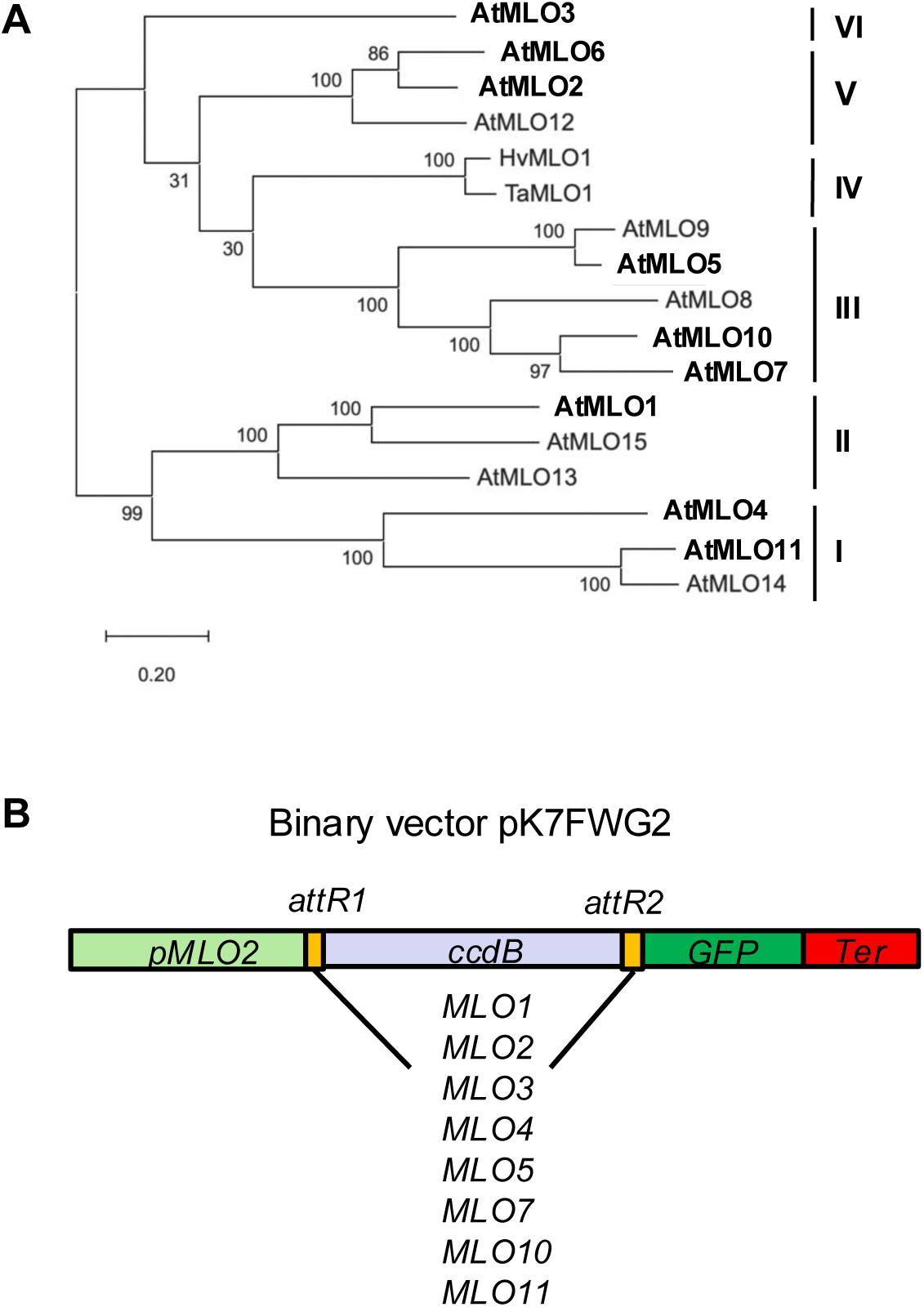
Selection and cloning of seven *MLO* family members from five different clades for ectopic expression in leaves of *eps3m* by the *MLO2* promoter. **(A)** A phylogenetic tree of the Arabidopsis MLO family plus barley and wheat clade IV MLO1 constructed based on deduced amino acid sequences using MEGA12 [Kumar S., Stecher G., Suleski M., Sanderford M., Sharma S., and Tamura K. (2024). Molecular Evolutionary Genetics Analysis Version 12 for adaptive and green computing. Molecular Biology and Evolution 41:1-9]. Bold-faced are MLO family members (belonging to different clades) that were subjected to expression and localization analyses. **(B)** Schematic showing the binary vector for expressing the eight indicated *MLO* genes from the *MLO2* promoter.

**Supplementary Figure S9.**
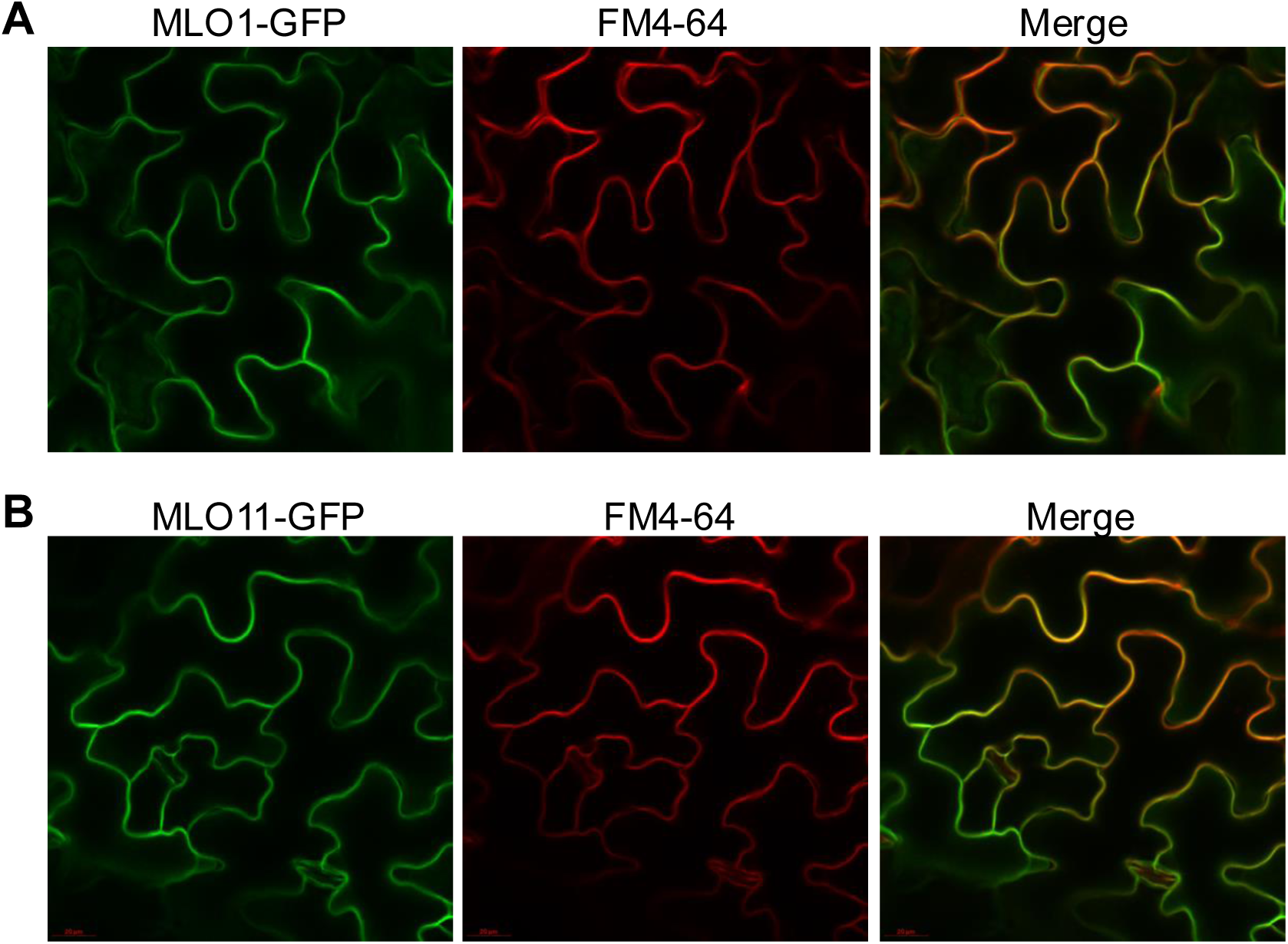
MLO1-GFP and MLO11-GFP are localized to the plasma membrane. Leaves of *eps3m* transgenic plants expressing MLO1-GFP (**A**) or MLO11-GFP (**B**) from the *MLO2* promoter were stained with 20 µM FM4-64 for 15 min before subjected to confocal imaging. Shown are representative z-stack confocal images projected from 3-5 thin optical sections.

**Supplementary Figure S10.**
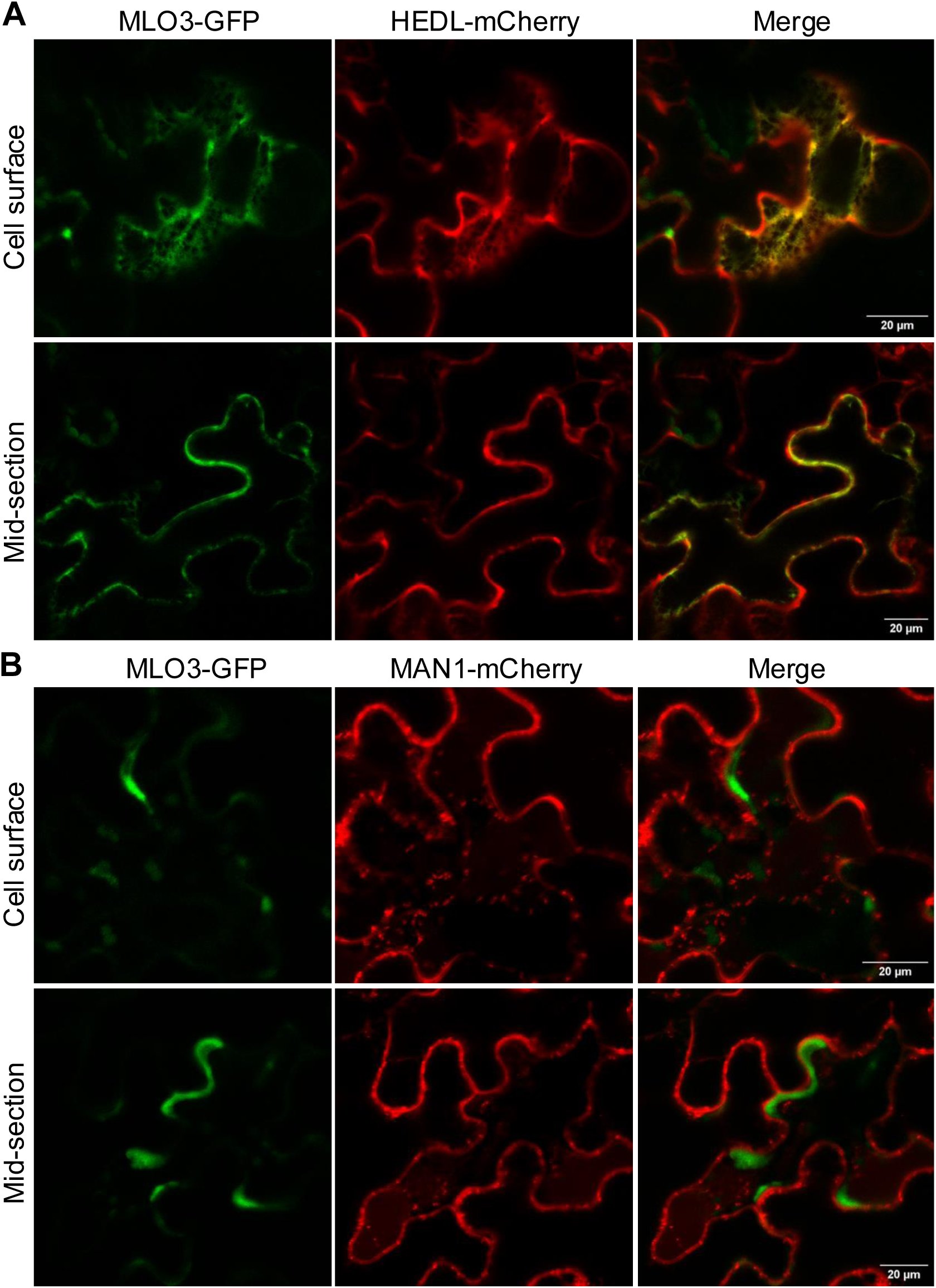
MLO3-GFP exhibits ER-localization in leaf epidermal cells of *N. benthamiana*. Agrobacterium cells harboring *pMLO2::MLO3-GFP* were mixed in equal concentration (OD_600_=0.5) with those harboring the ER marker *35S::HEDL-mCherry* (**A**) or the Golgi marker *35S::Man1-mCherry* (**B**). The mixtures were infiltrated into leaves of *N. benthamiana*. Confocal images were acquired at 2 days after agroinfiltration. Shown are Z-stack projections of 3-5 optical sections.

**Supplementary Figure S11.**
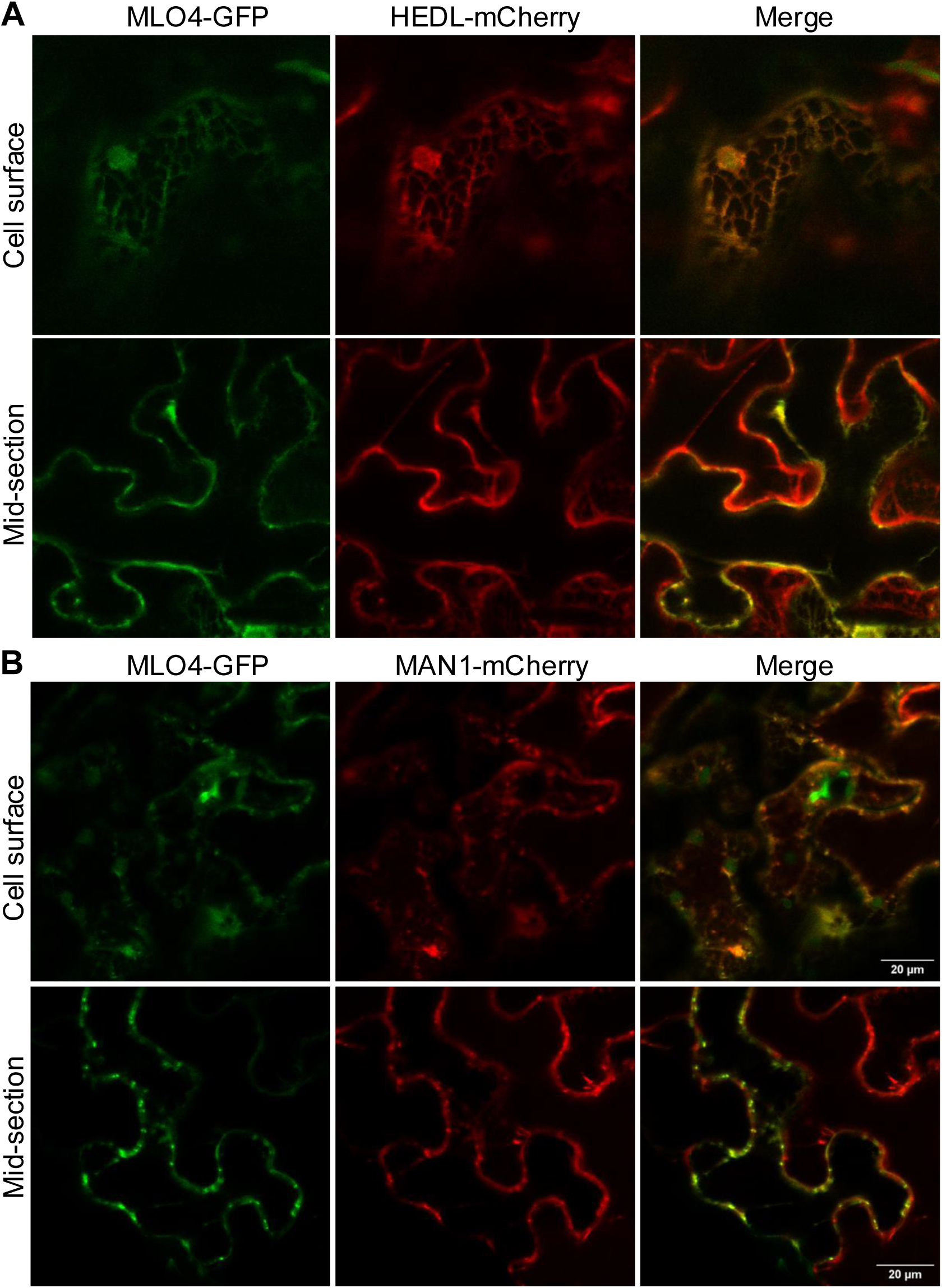
MLO4-GFP exhibits partial ER- and partial Golgi-localization in leaf epidermal cells of *N. benthamiana*. Agrobacterium cells harboring *pMLO2::MLO4-GFP* were mixed in equal concentration (OD_600_=0.5) with those harboring the ER marker *35S::HEDL-mCherry* (A) or the Golgi marker *35S::Man1-mCherry* (B). The mixtures were infiltrated into leaves of *N. benthamiana*. Confocal images were acquired at 2 days after agroinfiltration. Shown are Z-stack projections of 3-5 optical sections.

**Supplementary Figure S12.**
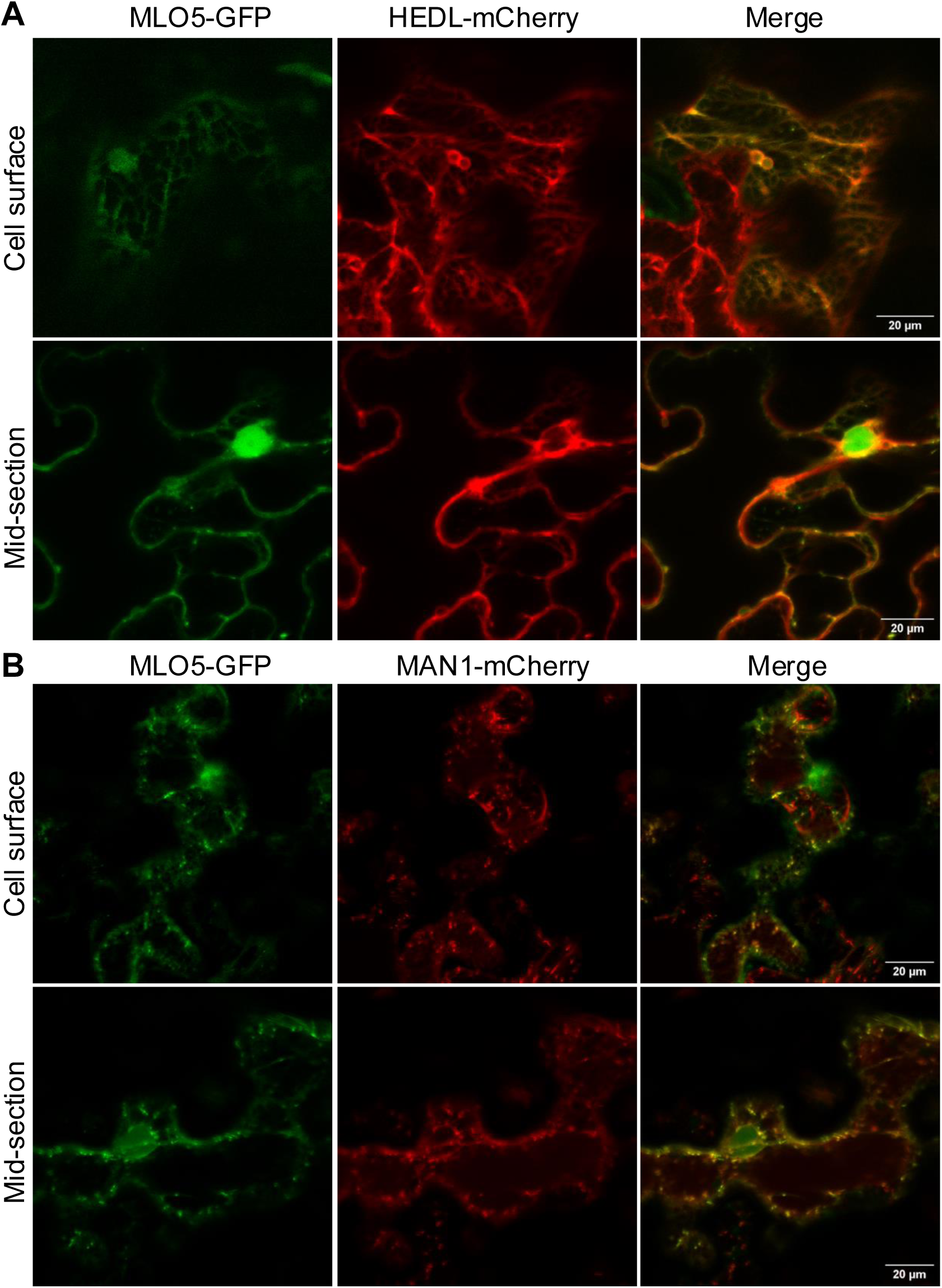
MLO5-GFP exhibits partial ER- and partial Golgi-localization in leaf epidermal cells of *N. benthamiana*. Agrobacterium cells harboring *pMLO2::MLO5-GFP* were mixed in equal concentration (OD_600_=0.5) with those harboring the ER marker *35S::HEDL-mCherry* (**A**) or the Golgi marker *35S::Man1-mCherry* (**B**). The mixtures were infiltrated into leaves of *N. benthamiana*. Confocal images were acquired at 2 days after agroinfiltration. Shown are Z-stack projections of 3-5 optical sections.

**Supplementary Figure S13.**
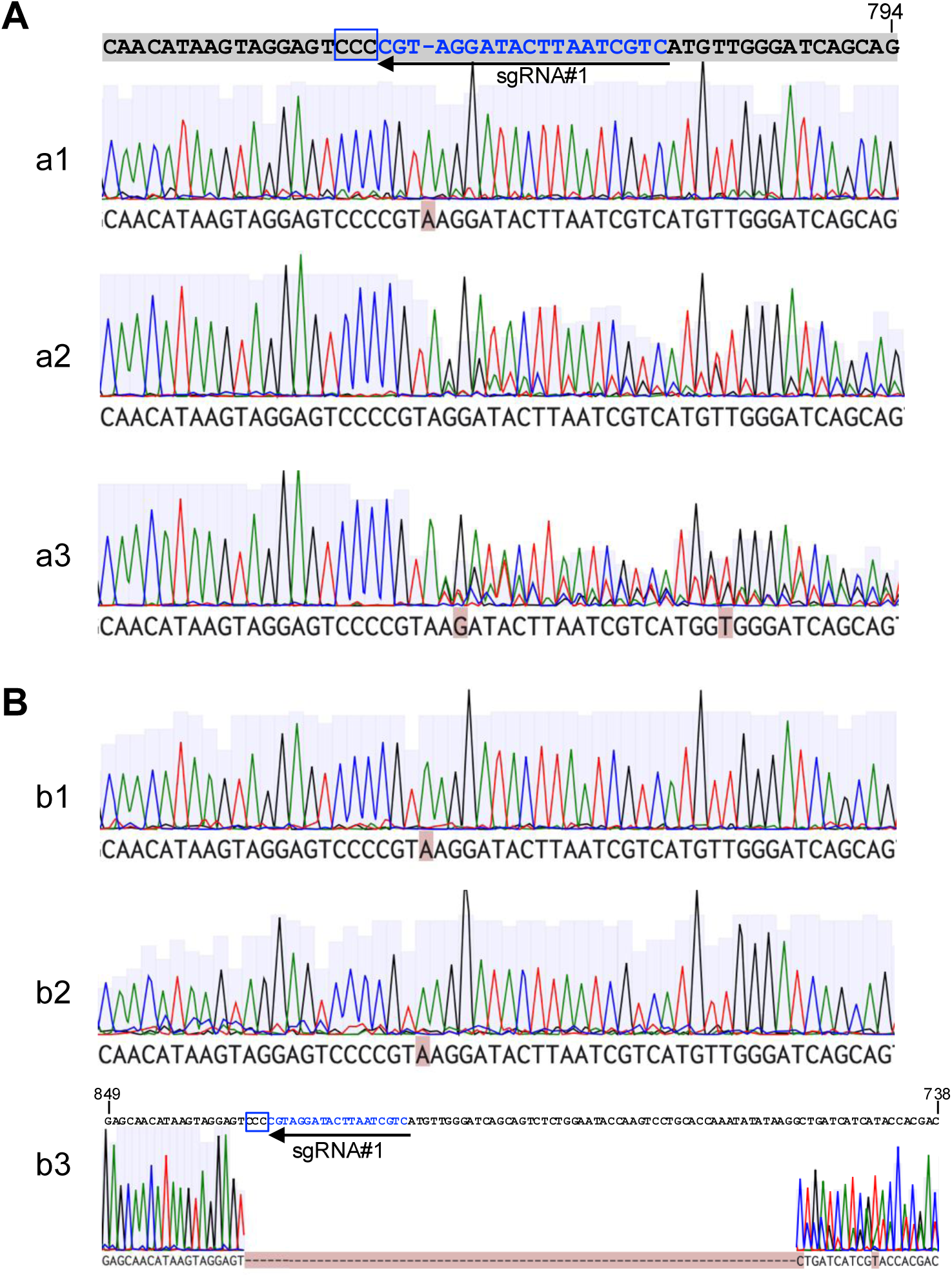
CRISPR-targeted mutagenesis of *FER*. The CRISPR construct was introduced in *eps* (A) and *eps3m* expressing MLO2-GFP and RPW8.2-RFP (B). **(A)** Sanger sequencing chromatograms of the sgRNA target regions of three independent T1 lines with a compact rosette in the *eds1/pad4/sid2* (*eps*) triple mutant background. **(B)** Sanger sequencing chromatograms of the sgRNA target regions of three independent T1 lines with a compact rosette in the background of *eds1/pad4/sid2/mlo2/mlo6/mlo12* (*eps3m*) plants transgenic for *pMLO2::MLO2-GFP* and *pRPW8.2::RPW8.2-RFP*.

**Supplementary Figure S14.**
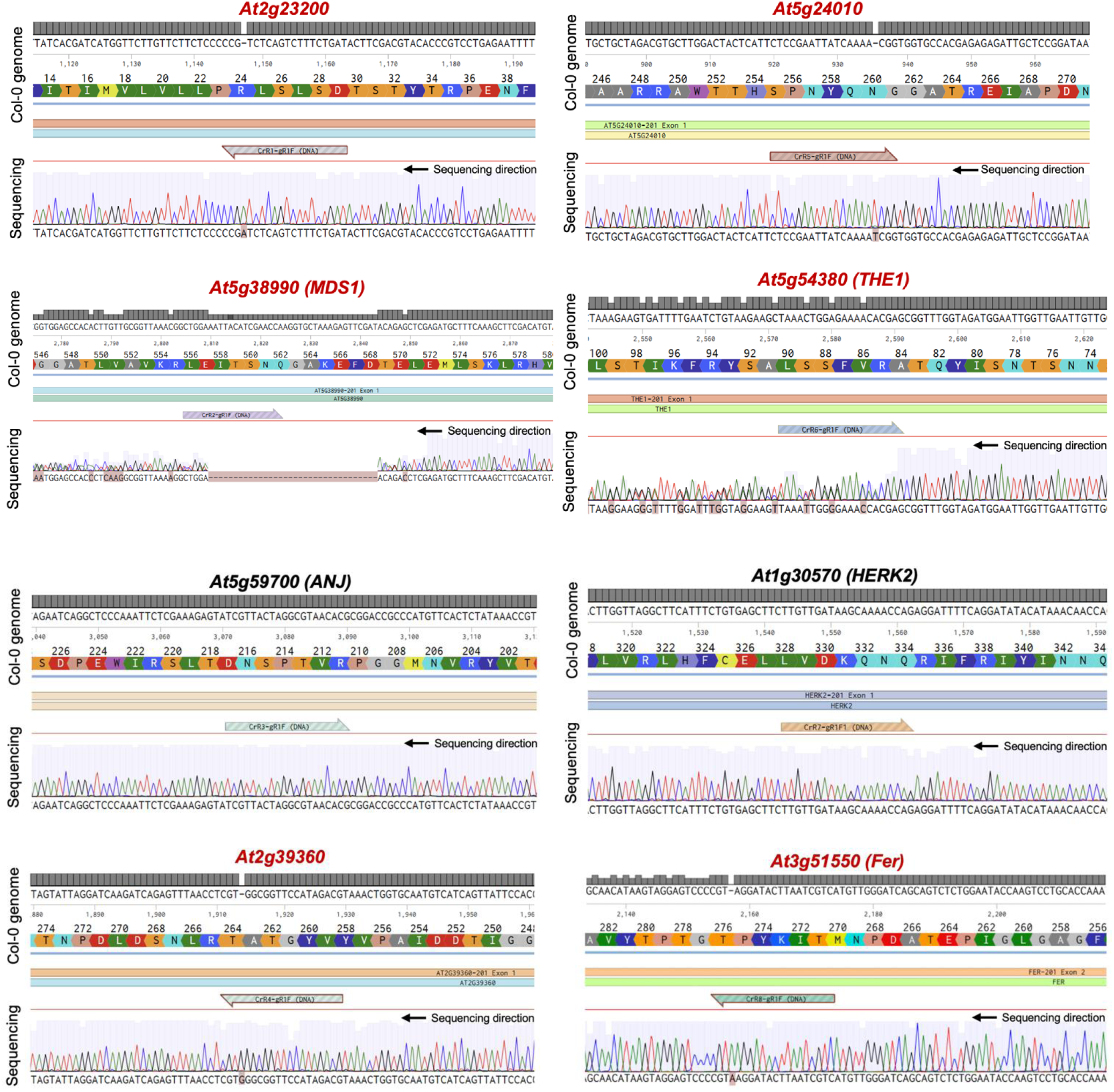
Multiplexed CRISPR targeting eight *CrRLK1L* family members. The multiplexed CRISPR construct was introduced into *eps3m* expressing MLO2-GFP and RPW8.2-RFP. Shown are the sequence alignments between the wild-type (Col-0) sequence and the Sanger sequencing chromatograms of the sgRNA-targeting regions of eight *CrRLK1L* genes in one mutant line (e2) exhibiting compact rosette and susceptibility phenotypes. Notably, six (highlighted in red) of the eight targeted *CrRLK1L* genes contain disruptive indels.

**Supplementary Figure S15.**
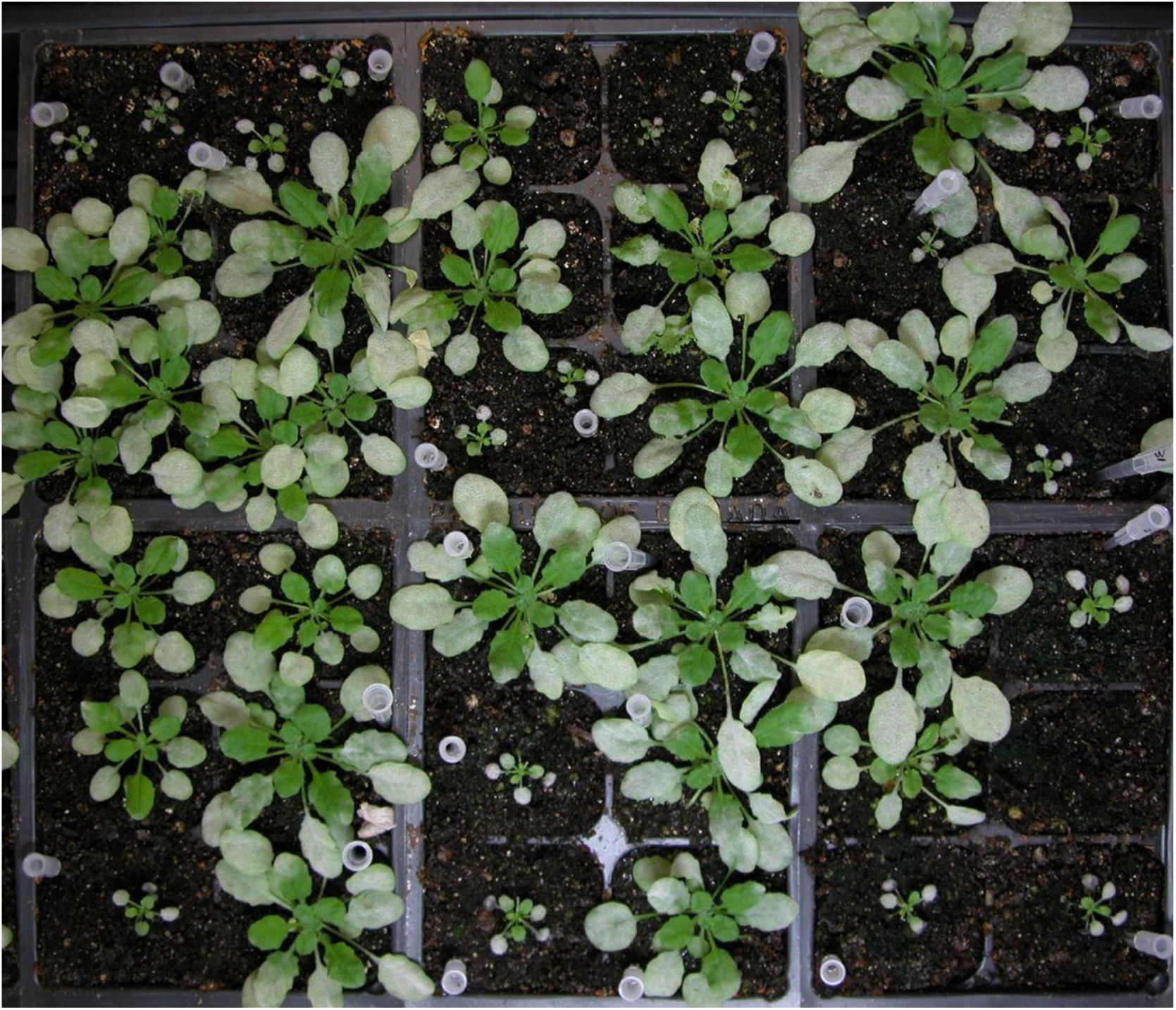
CRISPR-targeted mutagenesis of *PIP5K1* and *PIP5K2*. Shown are 42 T1 plants transgenic for a CRISPR construct targeting *PIP5K1* and *PIP5K2* in *eps3m* expressing MLO2-GFP and RPW8.2-RFP at 10 dpi with *Gc* UCSC1. Note, 17 of the 42 T1 plants exhibited greatly reduced stature and susceptibility to *Gc* UCSC1.

**Supplementary Table S1.**
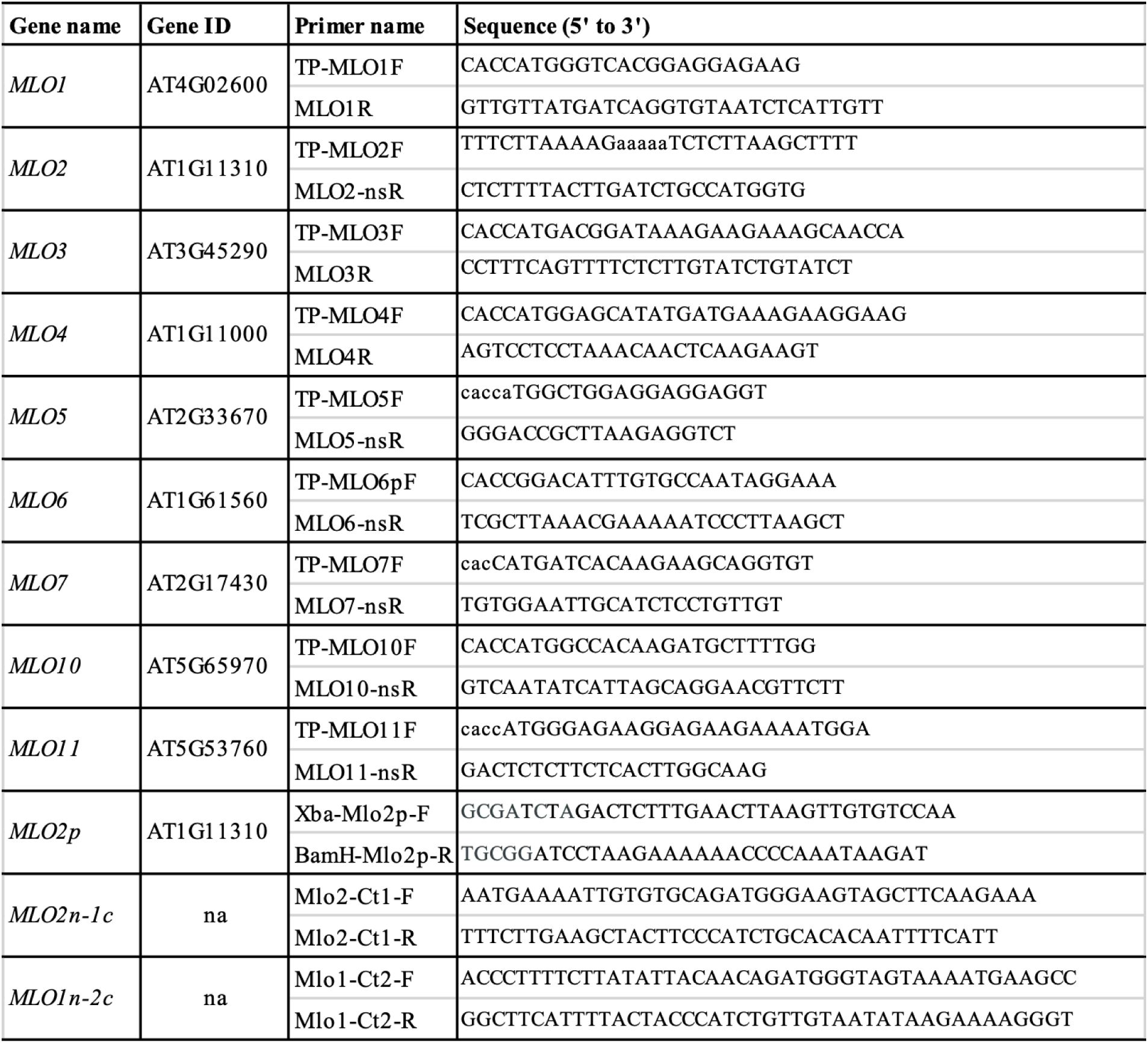
Primers used for cloning MLO genes.

**Supplementary Table S2.**
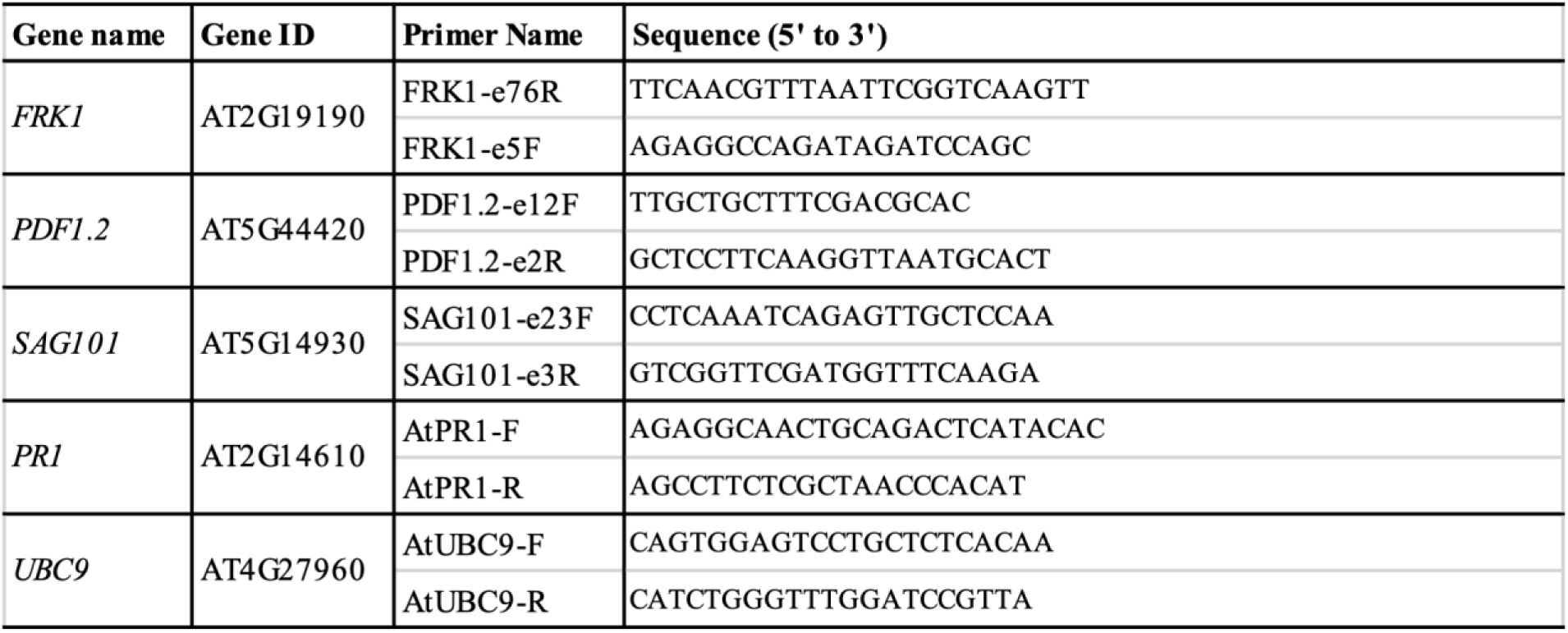
Primers used for gRT-qPCR.

**Supplementary Table S3.**
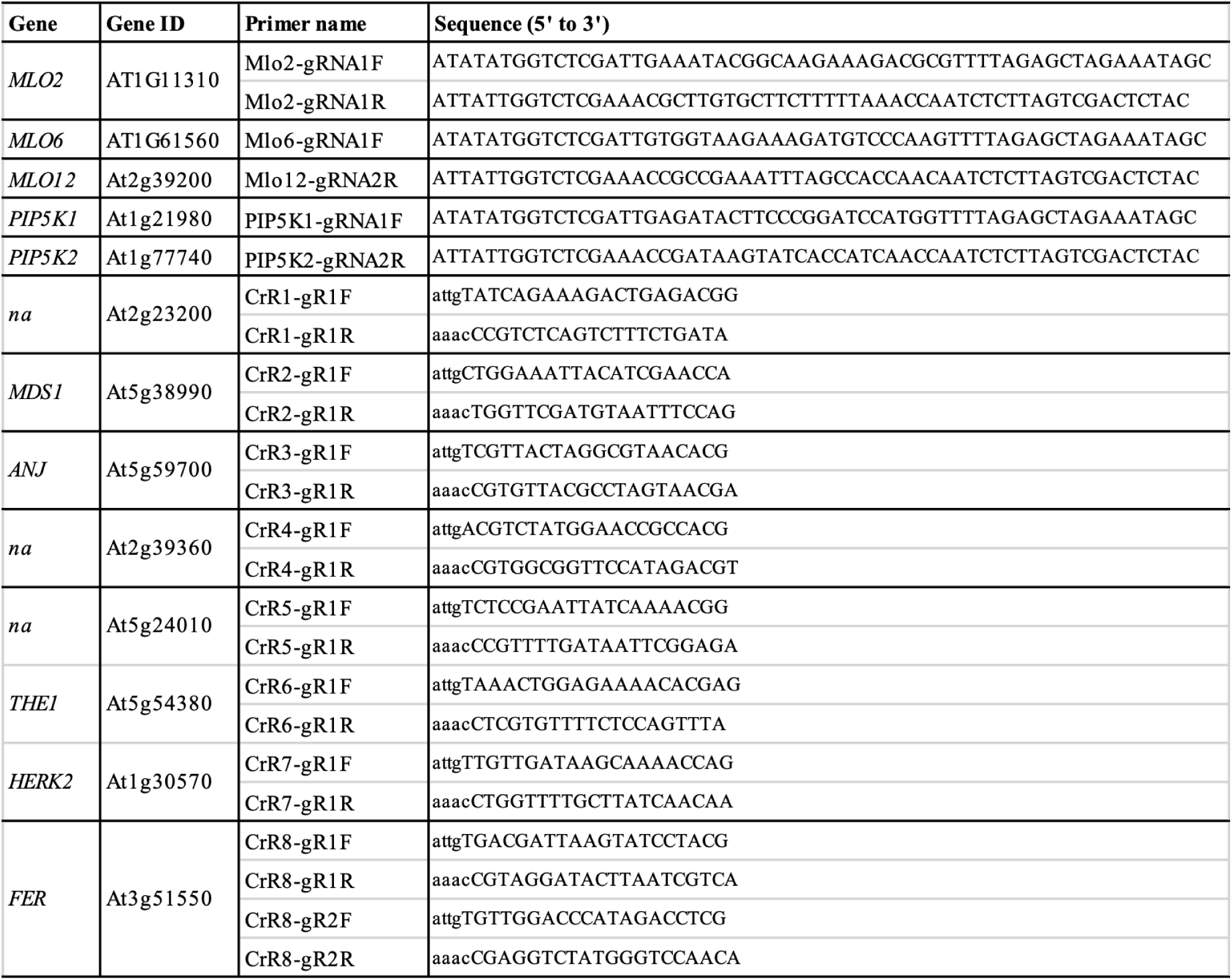
Small guide RNAs used for CRISPR/Cas9-targeted mutagenesis.

**Supplementary Table S4.**
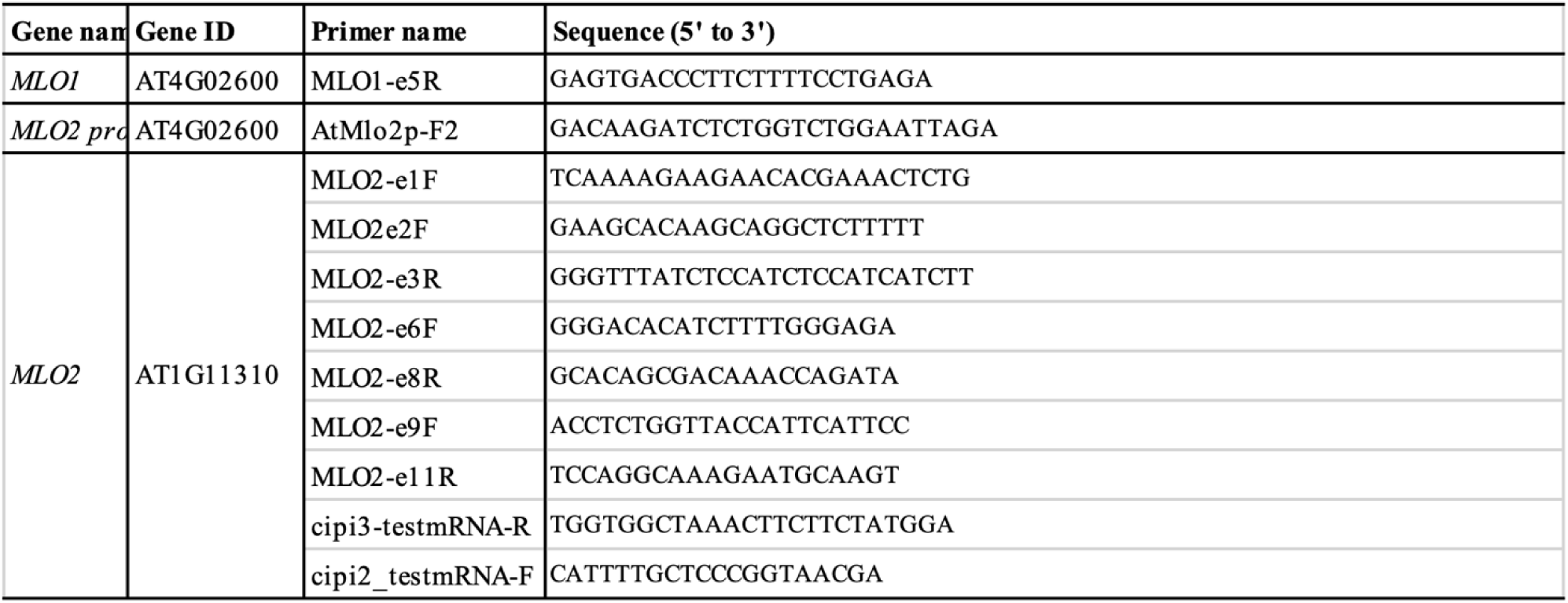
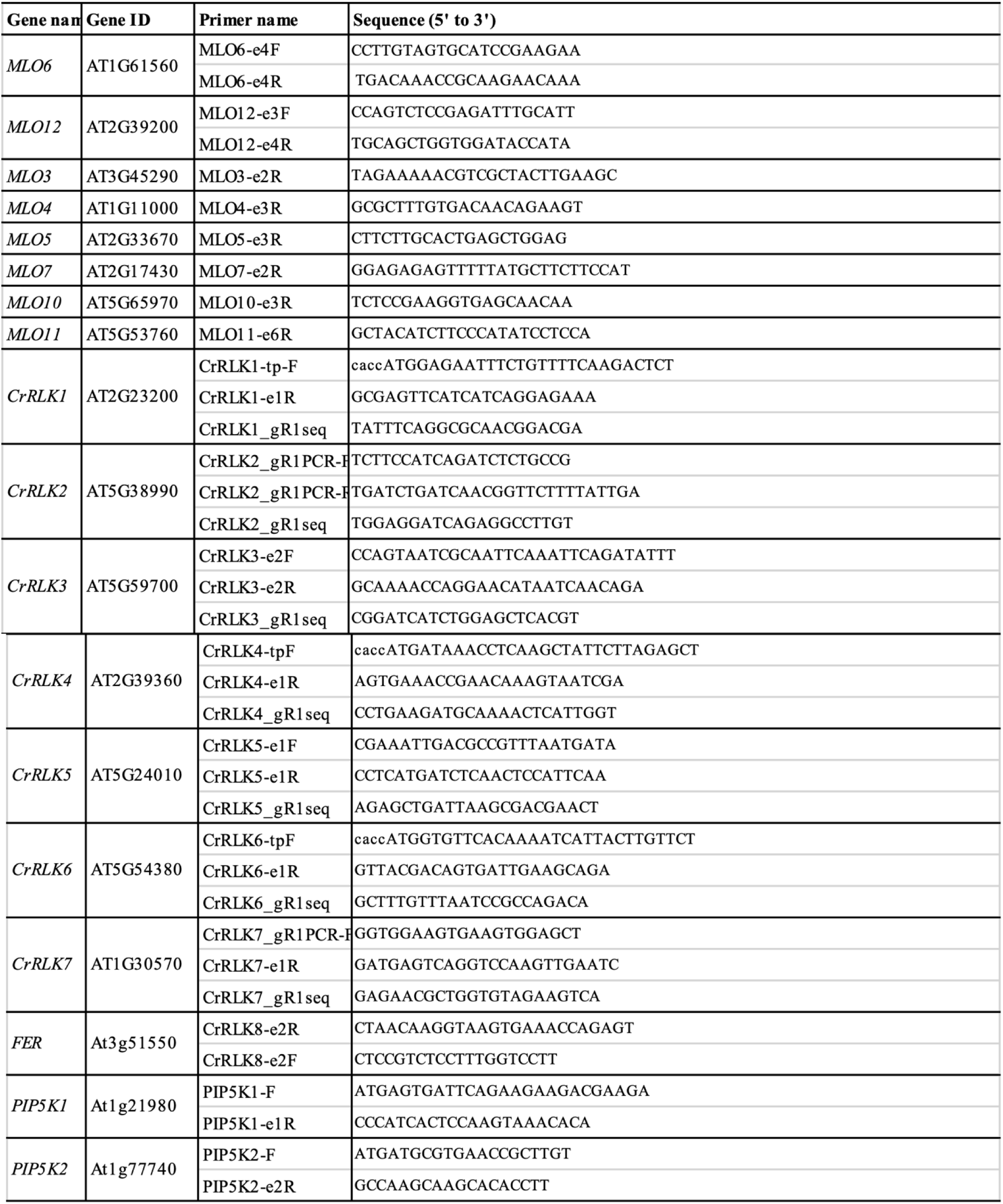
Primers used for genotyping or sequencing.

## ACKNOWLEDGMENTS

We thank Frank Coker and Caroline Hooks for their assistance in maintenance of plant growth facility a critically reading the article. This work was supported the National Sciences Foundation (IOS-1901566; IOS-2224203) to S.X.

## AUTHOR CONTRIBUTIONS

S.X., C-I.W., and Q.Z. designed and initiated the project; Q.Z. identified the *mlo2* mutants, D.B., Q.Z., S.X., Y.W., P.L., M.P., C.Z., A.H., J.Z., performed various experiments; R.P. and S.K. provided research materials and improve the manuscript; L.S. and P.H. helped with mutant characterization; D.B., Q.Z. and S.X. analyzed data and wrote the manuscript with help from other coauthors.

